# LARP1 is a major phosphorylation substrate of mTORC1

**DOI:** 10.1101/491274

**Authors:** Bruno D. Fonseca, Jian-Jun Jia, Anne K. Hollensen, Roberta Pointet, Huy-Dung Hoang, Marius R. Niklaus, Izabella A. Pena, Roni M. Lahr, Ewan M. Smith, Jaclyn Hearnden, Xu-Dong Wang, An-Dao Yang, Giovanna Celucci, Tyson E. Graber, Christopher Dajadian, Yonghao Yu, Christian K. Damgaard, Andrea J. Berman, Tommy Alain

**Affiliations:** Children’s Hospital of Eastern Ontario (CHEO) Research Institute, Department of Biochemistry, Microbiology and Immunology, University of Ottawa, Ottawa, Canada; Department of Molecular Biology and Genetics, Aarhus University, Aarhus, Denmark; Department of Biological Sciences, University of Pittsburgh, Pittsburgh, Pennsylvania, United States; Cancer Research UK Beatson Institute, Glasgow, United Kingdom; Department of Biochemistry, Rosalind and Morris Goodman Cancer Centre, McGill University, Montréal, Canada; Department of Biochemistry, UT Southwestern Medical Center, Dallas, Texas, United States

## Abstract

The mammalian target of rapamycin complex 1 (mTORC1) controls critical cellular functions such as protein synthesis, lipid metabolism, protein turnover and ribosome biogenesis through the phosphorylation of multiple substrates. In this study, we examined the phosphorylation of a recently identified target of mTORC1: La-related protein 1 (LARP1), a member of the LARP superfamily. Previously, we and others have shown that LARP1 plays an important role in repressing TOP mRNA translation downstream of mTORC1. LARP1 binds the 7-methylguanosine triphosphate (m^7^Gppp) cap moiety and the adjacent 5’terminal oligopyrimidine (5’TOP) motif of TOP mRNAs, thus impeding the assembly of the eIF4F complex on these transcripts. mTORC1 plays a critical role in the control of TOP mRNA translation *via* LARP1 but the precise mechanism by which this occurs is incompletely understood. The data described herein help to elucidate this process. Specifically, it show that: (i) mTORC1 interacts with LARP1, but not other LARP superfamily members, *via* the C-terminal region that comprises the DM15 domain, (ii) mTORC1 pathway controls the phosphorylation of multiple (up to 26) serine and threonine residues on LARP1 *in vivo*, (iii) mTORC1 regulates the binding of LARP1 to TOP mRNAs and (iv) phosphorylation of S689 by mTORC1 is particularly important for the association of the DM15 domain of LARP1 with the 5’UTR of RPS6 TOP mRNA. These data reveal LARP1 as a major substrate of mTORC1.

## Introduction

mTORC1 plays a fundamental role in the control of mRNA translation (1–5), particularly that of a class of transcripts known as TOP mRNAs, so-called because they bear a 5’terminal oligopyrimidine sequence immediately downstream of the N^7^-methyl guanosine triphosphate mRNA cap (6,7). The translation of TOP mRNAs is exquisitely sensitive to mTORC1 inhibition achieved by pharmacological agents (8–10), or nutrient- (8,11), growth factor- (12), hormone- (13) and oxygen-deprivation (14). Remarkably, TOP mRNAs are the class of transcripts that is most sensitive to translation repression upon mTOR inhibition (15–17).

The existence of a repressor of TOP mRNA translation has been recognized since 1999 (18), but the identity of said repressor and the precise mechanism by which mTORC1 controlled its activity remained unclear (6). Recent work has connected a novel signaling axis (the mTORC1-LARP1 signaling axis) to the control of TOP mRNA translation (19,20). La-related protein 1 (LARP1), an RNA-binding protein, represses the translation of TOP mRNAs downstream of mTORC1 (19). Our initial data suggested that it does so by competing with the eIF4F complex for binding to TOP mRNAs (19) but the exact mechanism by which this occurs was, until recently, incompletely understood. Recent structural (21,22) and biochemical (23) data revealed that LARP1 interacts with the m^7^Gppp cap and the adjacent 5’TOP motif *via* its conserved carboxy-terminus DM15 domain (21,22). In doing so, LARP1 effectively displaces eIF4E from the m^7^Gppp cap of TOP mRNAs and precludes the association of eIF4G1 with TOP mRNAs (22), thus blocking TOP mRNA translation (23,24).

How does mTORC1 dictate the inhibitory activity of LARP1? Typically, mTORC1 modulates the activity of its downstream targets through multisite phosphorylation of key serine and threonine residues. For instance, mTORC1 catalyzes the phosphorylation^1^ of multiple residues on ribosomal protein S6 kinases (S6Ks) (26–31), eukaryotic initiation factor 4E-binding proteins (4E-BPs) (32–43) and proline-rich AKT substrate 40kDa (PRAS40) (44,45), a less well-characterized substrate of mTORC1 (25). 4E-BPs (of which there are three homologs in mammals: 4E-BP1, 4E-BP2 and 4E-BP3) and S6Ks (S6K1 and S6K2) are the most intensively studied direct mTORC1 substrates; accordingly, these two targets are frequently referred to as the most important effectors of mTORC1 in mRNA translation (46). Two authoritative phosphoproteome studies (47,48) coupled the use of mTOR-specific pharmacological agents (rapamycin and torin1/Ku-0063794) to the power of liquid chromatography tandem mass spectrometry (LC-MS/MS) to reveal that, in addition to the well-characterized 4E-BPs and S6Ks, the mTORC1 pathway modulates the phosphorylation (either directly or indirectly by way of activation of downstream kinases) of thousands of presently uncharacterized mTORC1 substrates. LARP1 is one such new mTORC1 substrate (47,48). mTORC1 directly catalyzes the phosphorylation of LARP1 *in vitro* (49,50) but the significance of this phosphorylation event is presently unknown. In this study, we demonstrate that rapamycin alters the phosphorylation of LARP1 at multiple serine and threonine residues (and somewhat unexpectedly at tyrosine residues). Importantly, from a regulatory standpoint, our data show that phosphorylation of LARP1 residue S689 diminishes its RNA binding activity. This study provides the first evidence for a modulatory role for mTORC1-mediated LARP1 phosphorylation.

## Results

### mTORC1 interacts most strongly with LARP1 (among LARP superfamily members)

LARP1 belongs to the LARP superfamily (illustrated in **Fig. 1A**). In mammals, the LARP superfamily is comprised by 7 members: La also known as genuine La (the founding member of this superfamily), LARP1, LARP2 (also known as LARP1b), LARP4, LARP5 (also referred to as LARP4b), LARP6 and LARP7 (25,51,52). We have previously shown that LARP1 interacts with mTORC1 *via* the scaffolding protein RAPTOR (short for regulatory associated protein of mTOR), specifically through contact points with the RNC (RAPTOR N-terminal conserved) region and the WD40 (tryptophan/aspartate repeats 40 amino acids-long) domain (19). Given the high level of sequence conservation between human LARP1 and human LARP2 proteins (60% identity and 73% similarity at amino acid level^2^), we asked whether LARP2 also interacted with mTORC1. HEK293T cells were transiently transfected with FLAG-tagged human LARP1, human LARP2 or a variant of human LARP2 that lacks the C-terminal region (human LARP2 ΔCT), and subsequently stimulated transfected cells with complete growth media (see experimental procedures section for details). Cells were then lysed and lysates subjected to immunoprecipitation with anti-FLAG tag antibody. FLAG-immunoprecipitates were probed for endogenous RAPTOR and endogenous mTOR proteins by SDS-PAGE/Western blot. The resulting data (shown in **Fig. 1B**) confirmed our earlier observation (19) that LARP1 (in this particular case exogenous LARP1) interacts strongly with both RAPTOR and mTOR; this interaction is visualized on both short and long Western blot exposures. By contrast, FLAG-LARP2 interacted only very weakly with mTORC1 (**Fig. 1B**); endogenous RAPTOR and mTOR bands are barely detectable on short exposure and can only be visualized by Western blot upon saturation of the signal (**Fig. 1B**). The FLAG-LARP2 ∆CT variant failed to interact with endogenous RAPTOR or mTOR altogether. These data indicate that LARP1 – but likely not LARP2 – is a direct target of mTORC1. We extended this analysis to other LARP superfamily members (**Fig. 1C**). Similar to LARP2, La, LARP4, LARP5, LARP6 and LARP7 all failed to bind RAPTOR (**Fig. 1C**). Collectively, these data show that LARP1 is the sole member of this superfamily of proteins that is capable of interacting <u>avidly</u> with mTORC1.

**Figure 1.**
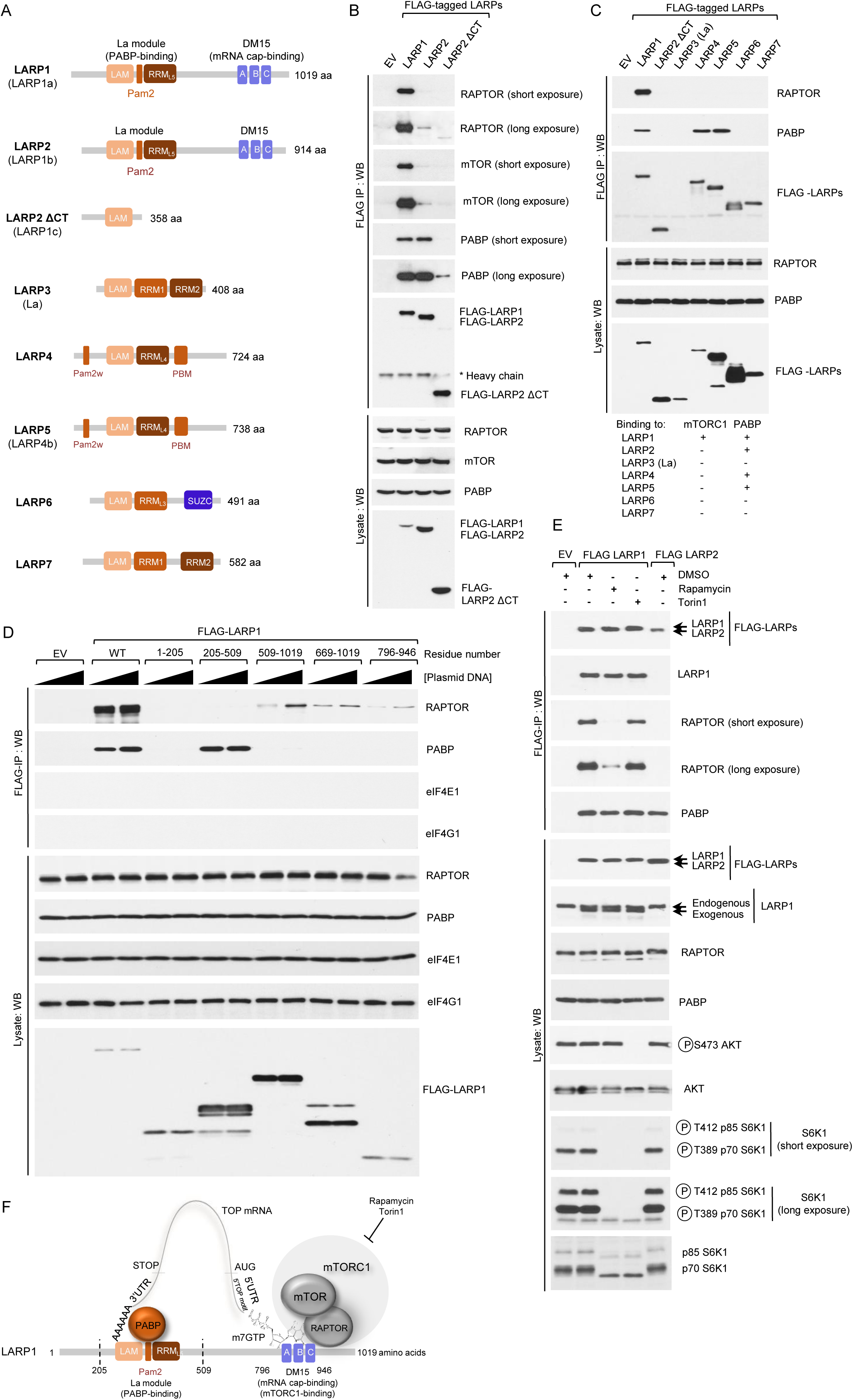
LARP1 and LARP2 bind PABP via the La module, while only LARP1 binds mTORC1 via the DM15 region in a rapamycin-sensitive manner. (A) Diagrammatic representation of the human LARP superfamily. (B) LARP1 and LARP2 interact with PABP, while only LARP1 binds mTORC1. HEK293T cells were transiently transfected with empty vector (EV), FLAG-tagged LARP1, –LARP2 or a variant of LARP2 that lacks the C-terminal region (-LARP2 ΔCT). Cells were stimulated with full growth media for 3 h and lysed in CHAPS buffer (see *Experimental Procedures* section). Lysates were subjected to immunoprecipitation and analyzed by SDS-PAGE/western blot with the indicated antibodies. (C) LARP1, LARP2, LARP4 and LARP5 interact with PABP. HEK293T cells were transfected and treated as described in (B) and lysates subjected to immunoprecipitation with anti-FLAG antibody. (D) RAPTOR binds to the C-terminal region of LARP1 that comprises the DM15 domain, while PABP binds to the La module that comprises the previously characterized (19) Pam2 motif. LARP1 does not interact with eIF4G1 or eIF4E1. HEK293T cells were transfected with empty vector (EV), full length FLAG-tagged wildtype LARP1 (WT) or FLAG-tagged fragments of LARP1. Amino acid numbering indicated. Transfected cells were treated as described in (A) and lysed in CHAPS lysis buffer (see *Experimental Procedures* section). Lysates were subjected to immunoprecipitation with anti-FLAG antibody and immunoprecipitates and lysates analysed by SDS-PAGE/western blot with the indicated antibodies. (E) LARP1 binds to mTORC1 in a rapamycin-sensitive manner, while Torin1 has no discernible effect on this interaction. LARP2 does not bind to mTORC1, nor does it bind to LARP1. HEK293T cells were transfected with FLAG-tagged LARP1 or –LARP2. Where indicated cells were stimulated with full-growth media in the presence of 0.1% (v/v) DMSO (vehicle), 100 nM rapamycin or 300 nM torin1 for 3 h. Cells were lysed as described in (A-D) and lysates subjected to immunoprecipitation with anti-FLAG antibody. IPs and lysates were analyzed as described in (A-D). (F) Diagrammatic representation of the various domains in LARP1 and its interaction with mTORC1, PABP and TOP mRNAs. La module mediates PABP-binding while the DM15 mediates mRNA cap- and mTORC1-binding.

### The LARP superfamily members: LARP1, LARP2, LARP4 and LARP5 interact with PABP

The LARP superfamily is so-named because its members share an evolutionarily-conserved ancestral La module that encompasses the La motif (LaM) and a RNA-recognition motif (RRM) (**Supplemental Fig. 1**) that play important functions in RNA biology (reviewed in (51)). In addition to the shared La module domain, a number of LARPs display unique features such as the PAM2 motif that mediates binding to the *mademoiselle* (MLLE) motif in the poly(A)-binding protein (PABP) (53,54). LARP1 contains a *bona fide* canonical PAM2 motif – nudged between the La motif and RRM-like 5 (RRM_L5_) (**Fig. 1A**) – that mediates binding to PABP (19). The PAM2 motif is also conserved in LARP2, a close homolog of LARP1 (**Fig. 1A and Supplemental Fig. 1**); nothing is known about the function of LARP2. With these facts in mind, we tested whether LARP2 also binds PABP. To this end, we transiently overexpressed C-terminally tagged full-length human LARP1, human LARP2 or human LARP2 ∆CT followed by immunoprecipitation with a FLAG-specific antibody in the presence of ribonuclease A (RNase A) to exclude RNA-mediated protein-protein interactions (**Fig. 1B**). As expected from the PAM2 sequence conservation (**Supplemental Fig. 2**), both full length-LARP1 and -LARP2 interacted with endogenous PABP to similar extent (**Fig. 1B**). LARP2 ∆CT which lacks both the DM15, the RRM_L5_ and the Pam2 motif lost most of its capability of binding to PABP (**Fig. 1B**); a small amount of PABP bound to LARP2 ∆CT can be detected on the Western blot long exposure, indicating that regions N-terminal to the Pam2 motif in LARP2 likely also contribute, albeit weakly, to PABP binding. LARP4 (55) and LARP5 (56) have been previously demonstrated to also interact with PABP. They do so through two distinct regions: (i) *via* an atypical PAM2w motif that contains a tryptophan in place of the critical phenylalanine located in the N-terminal region of LARP4 and LARP5 (55) and (ii) *via* a PABP-binding motif (PBM) positioned C-terminally of the RRM_L4_ (56) (**Fig. 1A**). Having established that LARP1 (19) and LARP2 interact with PABP (**Fig. 1B**), next we sought to extend the PABP-interaction analysis to all human LARPs (**Fig. 1C**). Consistent with earlier reports (55,56), LARP4 and LARP5 – but not La, LARP6, or LARP7 – also interact with PABP (**Fig. 1C**).

### LARP1 interacts with PABP via the mid-region that comprises the La module and with mTORC1 via the carboxy-terminal region that comprises the DM15 motif

Next, we assessed which region(s) of LARP1 mediates the interactions with PABP and mTORC1. To this end, we overexpressed FLAG-tagged full-length human LARP1 or fragments of human LARP1 covering various regions of the protein (**Fig. 1D**). Full-length LARP1 efficiently co-precipitated endogenous RAPTOR and endogenous PABP. By contrast, fragment 1–205 (comprising the N-terminal region of LARP1 did not co-precipitate RAPTOR or PABP; fragment 205–509 (comprising the mid-region of LARP1) efficiently co-precipitated PABP but not RAPTOR. Conversely, the C-terminal fragments weakly co-precipitated RAPTOR but not PABP (**Fig. 1B**). Together, these data indicate that LARP1 interacts with PABP *via* its mid-region that spans the entire La module (consistent with earlier data on PAM2 (19)) and with RAPTOR through the C-terminal region that comprises the DM15 motif. Importantly, LARP1 did not interact with eukaryotic initiation factors 4E1 (eIF4E1) or 4G1 (eIF4G1), in contrast to earlier reports that suggested that LARP1 binds the eIF4F complex (20).

### Rapamycin disrupts the binding of LARP1 to mTORC1 but not that of LARP1 to PABP, while torin1 affects neither interaction

Next, we tested whether pharmacological inhibition of mTORC1 plays a role in regulating the interaction between LARP1 and mTORC1 or LARP1 and PABP. HEK293T cells were transiently transfected with FLAG-LARP1 or -LARP2 (used as a negative control for the interaction with mTORC1 and as a positive control for the interaction with PABP) and the cells were then stimulated with full growth media (containing serum, amino acids and glucose) in the presence of DMSO (vehicle), rapamycin or torin1; cells were then lysed and protein lysates subjected to immunoprecipitation with anti-FLAG antibody. FLAG-immunoprecipitates were analyzed by SDS-PAGE/Western blot with the indicated antibodies. As shown in **Fig. 1E**, rapamycin efficiently disrupted the interaction between LARP1 and RAPTOR but not that of LARP1 with PABP, indicating that allosteric inhibition of mTORC1 interferes with the LARP1/RAPTOR interaction but not of that of LARP1 with PABP (consistent with an earlier report (19)). In parallel, we also tested the effect of torin1 on the interaction between mTORC1 and LARP1. Surprisingly, torin1 had only a minor effect (**Fig. 1E**), suggesting that the rapamycin-mediated disruption of LARP1-RAPTOR interaction likely arises from an allosteric effect of rapamycin on the stability of the mTORC1 complex (57), rather than from a shut-off of mTOR catalytic activity.

### Characterization of the binding of LARP1 to RAPTOR

RAPTOR, an mTORC1-specific component (58–60), functions as scaffolding protein that bridges the interaction between mTOR and its downstream substrates. RAPTOR interacts with its binding partners (4E-BPs, S6Ks, PRAS40 and HIF1a) via a TOR signaling (TOS) motif (61,62). The TOS motif is characterized by a short peptide sequence (5 amino acids long) in which the first residue is invariably a phenylanine (61,62) (**Fig. 2A**); the remaining four residues accommodate a number of amino acid combinations, and a consensus sequence for the TOS motif has been loosely defined as (F_1_-(DEVMQPRA)_2_-(IMLE)_3_-(DVEL)_4_-(ELIQDA)_5_) (63) (**Fig. 2A**). Each of the following targets of mTORC1: 4E-BPs, S6Ks, PRAS40 and HIF1a contains a *bona fide* TOS motif that is essential for their interaction with mTORC1 (44,45,61,62,64) (**Fig. 2A**). Notably, the conserved phenylalanine at position 1 on the TOS motif of each of these proteins is essential for their interaction with RAPTOR. The TOS motif sequence in PRAS40 is defined by the amino acid combination FVMDE (44,45) (**Fig. 2A**) and the mode of binding of this sequence to RAPTOR has been recently characterized in atomic detail (65) (**Fig. 2B**). A single amino acid substitution (phenylalanine to alanine) within position +1 of the TOS motif of PRAS40 (FVMDE to AVMDE) abolishes its binding to RAPTOR (**Fig. 2C**) (44,45). Double mutation of the methionine and aspartate residues (at +3 and +4 positions respectively) (FVMDE to FVAAE) had a similar disruptive effect on the interaction between PRAS40 and RAPTOR (**Fig. 2C**). Close inspection of LARP1 primary sequence revealed a putative TOS motif (F889RLDI, numbering according to the 1019 amino acid-long human LARP1 isoform) that satisfied the TOS consensus sequence described previously (63) (**Fig. 2D**). Furthermore, this sequence lies within the C-terminal region of LARP1 that mediates RAPTOR binding (**Fig. 1D**) and is not fully conserved in LARP2 (FRREI) (**Fig. 2E** and **Fig. 2F**), making it an ideal candidate for RAPTOR interaction. Indeed, deletion of the entire FRLDI region markedly reduces the interaction of LARP1 with RAPTOR and mTOR (**Fig. 2G**). However, substitution of the critical F889 for an alanine did not affect LARP1 binding to RAPTOR arguing against this sequence functioning as a TOS motif (**Fig. 2G**). Replacement of the leucine and aspartate residues (at positions +3 and +4, respectively), that in PRAS40 are required for RAPTOR binding (**Fig. 2C**), for arginine and glutamate respectively also did not affect LARP1 binding to RAPTOR (*i.e*. converting the putative FRLDI TOS motif in LARP1 to the FRREI LARP2 sequence does not affect binding to RAPTOR) (**Fig. 2H**). Since LARP2 does not bind RAPTOR, it follows that the FRLDI sequence in LARP1 is not essential for RAPTOR binding. To verify this further, we performed the converse experiment: we replaced the FRREI sequence in LARP2 for the FRLDI sequence (**Fig. 2H**). Introduction of the FRLDI sequence in LARP2 did not rescue the binding to RAPTOR. Collectively, these results demonstrate that this region of LARP1 is not a TOS motif in that it does not mediate RAPTOR binding. In light of these findings, we speculated that tertiary structure – rather than primary sequence – explained the interaction between LARP1 and RAPTOR. We next compared the structures of the DM15 domain of LARP1 (spanning residues 796–946 (21,22)) to those of RAPTOR in complex with the TOS motif of PRAS40 (65) and noticed structural similarities between the *bona fide* FVMDE in PRAS40 (**Fig. 2B**, inset) and the K924AKNLDI sequence in LARP1 (**Fig. 2I**). The two sequences display similar tertiary structures, despite the lack of primary sequence conservation. Moreover, the KAKNLDI sequence is located in a loop and is not conserved in LARP2 (**Fig. 2E**) – thus making it a prime candidate for RAPTOR binding. To test this hypothesis, we mutated the leucine and aspartate residues to alanine – that in PRAS40 abrogate RAPTOR binding (**Fig. 2A** and (45)) and tested the interaction of this LARP1 mutant (KAKNAAI) with mTORC1 (**Fig. 2J**). This double mutant retained full binding capacity to RAPTOR and mTOR, arguing against this sequence mediating LARP1 binding to mTORC1 (**Fig. 2J**). Consistent with this finding, substitution of this sequence for the QSKTQS found in LARP2 does not alter the binding of LARP1 to RAPTOR (**Fig. 2K**)). Together, these data indicate that neither FRLDI nor the KAKNLDI mediate RAPTOR binding.

**FIGURE 2.**
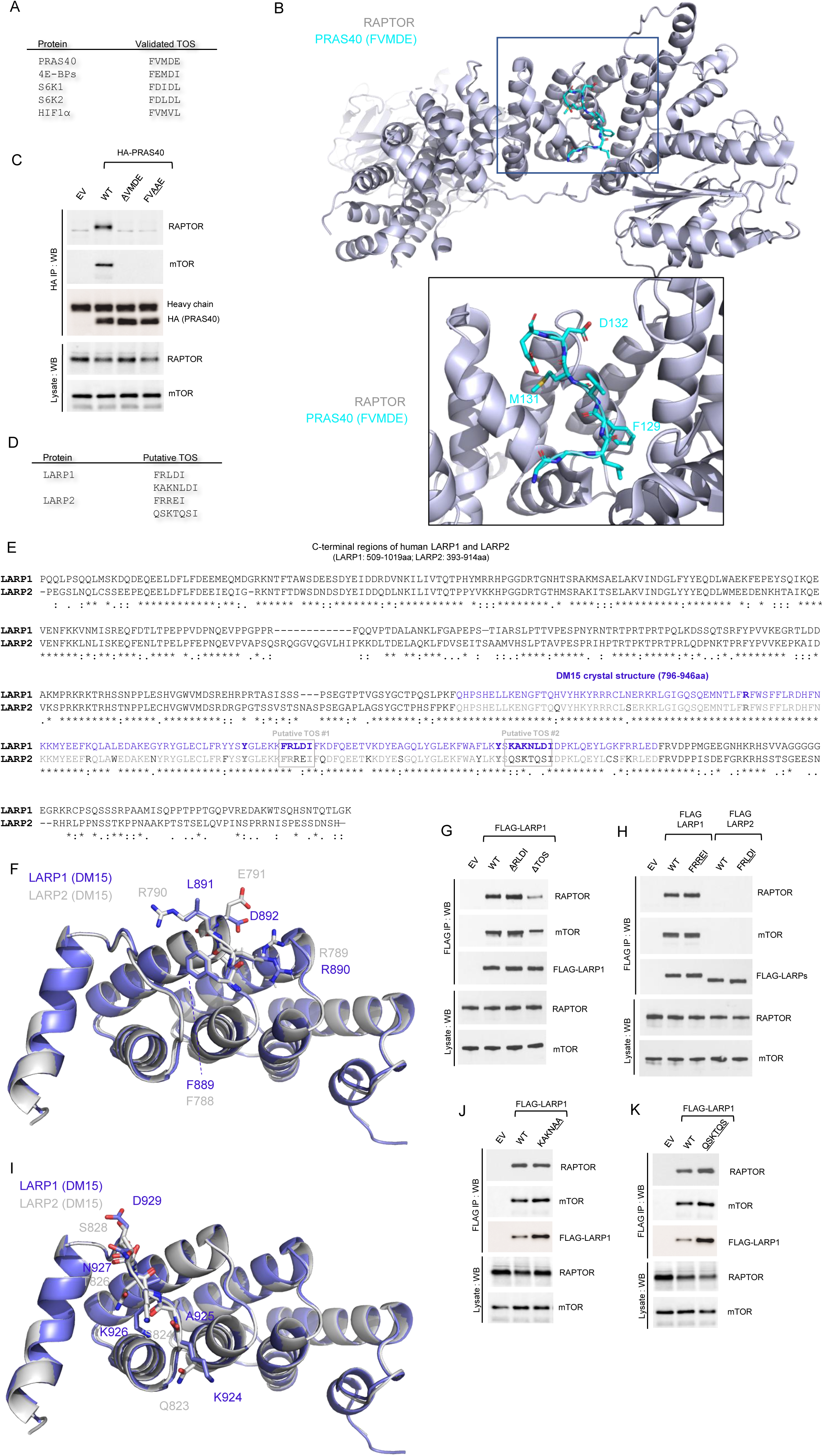
The FRLDI and KAKNLDI amino acid sequences in LARP1 do not mediate RAPTOR binding. (A) List of validated TOR signaling (TOS) motifs in well-characterized mTORC1 substrates. (B) Structure of PRAS40 TOS sequence in complex with RAPTOR. (C) F129, M131 and D132 residues are essential for PRAS40 binding to RAPTOR. (D) Putative TOS motifs in human LARP1 and human LARP2. (E) Alignment of C-terminal sections of human LARP1 and human LARP2. (F) Modeling of human LARP2 DM15 on solved human LARP1 structure displaying putative TOS motif #1 residues. (G-H) F889, L891 and D892 on LARP1 do not mediate RAPTOR binding. (I) Modeling of human LARP2 DM15 on solved human LARP1 structure displaying putative TOS motif #2 residues. (J-K) K924, A925, N927, L928, D929 on LARP1 do not mediate RAPTOR binding.

We have recently solved the structure of the DM15 domain of human LARP1 in complex with a segment of the 5’TOP motif of RPS6 mRNA as well as in complex with m^7^GTP and m^7^GpppC cap analogs (22). This published study revealed that the DM15 domain of LARP1 recognizes the mRNA cap moiety and the adjacent 5’TOP sequence of TOP mRNAs (as illustrated in **Fig. 3A**). We have shown that tyrosines 883 and 922 form a cation-pi stacking interaction with the N^7^-methylated guanosine moiety (akin to that of other cap binding proteins, *e.g*. eIF4E (66)), while arginine 840 forms a salt bridge with phosphate backbone of the 5’TOP sequence (22). The spatial arrangement of these residues is illustrated in **Fig. 3A** and the primary sequence conservation between human LARP1 and human LARP2 is shown in **Fig. 3B**. Previously, we have also noted that the binding of LARP1 to RAPTOR is enhanced upon degradation of RNA with RNase A (**Fig. 3C**) (19) leading us to hypothesize that RAPTOR and RNA (presumably TOP mRNAs) compete for LARP1 binding (19). In this context, we have previously shown (22) that R840 and Y883 are particularly important for LARP1 to bind the 5’end of RPS6 mRNA. With this in mind, we tested whether double mutation of R840 to a glutamate and Y883 to an alanine on LARP1 interfered with its association with RAPTOR. Our data showed that, indeed, simultaneous mutation of these two residues does abolish RAPTOR binding (**Fig. 3D**). We delved further into the contribution of each residue to RAPTOR binding by generating single mutations; mutation of R840 to a glutamate has the most dramatic effect on RAPTOR binding (**Fig. 3E**), almost as extensive as the double mutation. Mutation of Y883 to an alanine does not have a discernible effect on RAPTOR binding (**Fig. 3E**). These data indicate that charge reversal of R840 rather that mutation of Y883 is the key event for disruption of RAPTOR binding. This experiment identifies R840 as the first critical residue in LARP1 required for RAPTOR binding.

**FIGURE 3.**
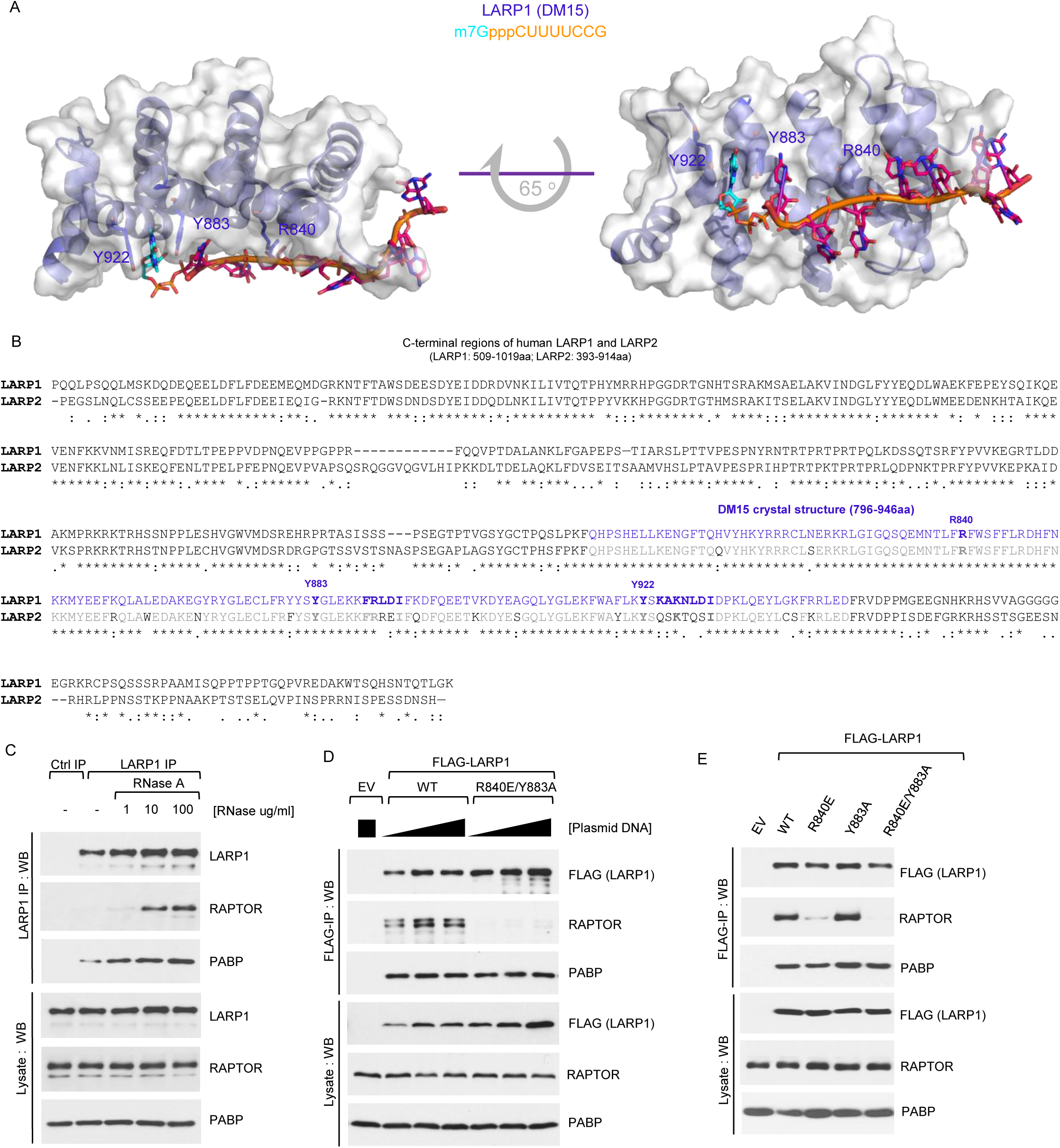
R840 on LARP1 is critical for its interaction with RAPTOR. (A) Molecular modeling of human LARP1 DM15 in complex of m7GpppCUUUUCCG sequence. (B) Human LARP1 and human LARP2 DM15 protein sequences alignment. (C) RNase A enhances the interaction between endogenous LARP1 and mTORC1. HEK293T cells were stimulated with complete media (experimental procedures) for 3 h and then lysed in CHAPS-based extraction buffer in the presence (of various concentrations) or absence of RNase A. Samples of lysates were subjected to immunoprecipitation with anti-LARP1 antibody and IPs/lysates analysed by SDS-PAGE/Western blot. (D) HEK293T cells were transiently transfected with empty vector (EV), human FLAG-tagged wildtype LARP1 (1019aa isoform) or the R840E/Y883A (REYA) double-mutant. Cells were stimulated and lysed as described in (C) in the presence of RNAse A. Lysates were subjected to immunoprecipitation with anti-FLAG antibody and analysed by SDS-PAGE/Western blot. (E) HEK293T cells were transiently transfected with empty vector (EV), human FLAG-tagged wildtype LARP1 (1019aa isoform) or the indicated mutants, stimulated, lysed and analysed as described in (D).

### The PT3K/AKT/mTORC1 signaling pathway plays a key role in regulating the binding of LARP1 to TOP mRNAs, while inhibition of S6K1 activity has a negligible effect on LARP1 association to TOP mRNAs

The PI3K/AKT/mTORC1 signaling pathway (depicted in **Fig. 4A**) plays a fundamental role in the control of TOP mRNA translation. This is documented in numerous pharmaco-genetic studies: inhibitors of this pathway such as rapamycin (8), LY294002 (a low-specificity PI3K inhibitor (67)) (12) and torin1 (a dual mTORC1/mTORC2 inhibitor) all reduce TOP mRNA translation (16), albeit to different degrees. Rapamycin partially reduces amino acid- (8) and serum-mediated (13) TOP mRNA translation, while torin1 has a more pronounced inhibitory effect on the translation of these transcripts (16). Equivalently, genetic ablation of key components of the PI3K/AKT/mTORC1 signaling pathway compromises TOP mRNA translation. For instance, genetic deletion of AKT1 (also referred to as PKBa, short for protein kinase B), impairs serum-mediated activation of TOP mRNA translation (12). Conversely, genetic deletion of the tuberous complex (TSC)-specific component TSC2 enhances basal translation of TOP mRNAs and renders it resistant to serum deprivation (13), indicating that signaling through this pathway is central to the control of TOP mRNA translation. Recently, we proposed that LARP1 functions as a critical repressor of TOP mRNA translation downstream of mTORC1 (19) based on several observations including that LARP1 physically interacts with mTORC1 and TOP mRNAs, and downregulation of its expression renders cells partially resistant to the inhibitory effects of rapamycin and torin1 on TOP mRNA translation (19). Notably, from a regulation standpoint, the binding of LARP1 to TOP mRNAs correlates with the activation status of mTORC1. Torin1, and to a lesser extent rapamycin, enhance the binding of endogenous LARP1 protein to TOP mRNAs (reciprocally these drugs also diminish the binding of endogenous eIF4G1 to these transcripts) (19). We considered it important to dissect further the contribution of each component of the PI3K/AKT/mTORC1 pathway (see **Fig. 4A**) to the binding of LARP1 to TOP mRNAs. To test this, HEK293T cells were acutely treated with mTOR inhibitors (rapamycin or torin1), an S6K1 inhibitor (PF4708671 (68)), an allosteric AKT inhibitor (MK2206 (69)) or a low-specificity PI3K inhibitor (LY294002 (67,70)) and lysates subjected to RNA immunoprecipitation (RNA-IP) using an antibody against endogenous LARP1 (Fig. 4B-E). The resulting data show that endogenous LARP1 preferentially interacts with highly abundant TOP mRNAs (e.g. RPS6 and RPL32, tested here) compared to other less abundant non-TOP mRNAs (e.g. lactate dehydrogenase A (LDHA) and beta-actin (β-actin)^3^ (**Fig. 4C**); this even upon normalization of LARP1-bound mRNA levels to their total abundance in the lysate/input (*cf*. **Fig. 4C** and **4E**)^4^. First, we noted that drug treatment has modest – yet, statistically significant – effects on mRNA steady-state levels. Of all the drugs, torin1 had the most drastic effect on mRNA steady-state levels – specifically, HEK293T cells treated with 300 nM torin1 for as short as 3 h appear to result in significantly (p-value <0.001) decreased mRNA RPS6, RPL32 and LDHA mRNAs association (**Fig. 4D**); this was not evident for β-actin mRNA (p-value is non-significant) possibly due to its intrinsic low abundance compared to the other mRNAs. Importantly, incubation of cells with rapamycin, torin1, MK2206, and LY294002 all enhanced the binding of LARP1 to all mRNAs tested including TOP (RPS6 and RPL32) and non-TOP (LDHA and β-actin) mRNAs (**Fig. 4C** and **Fig. 4E**)^4^. Rapamycin leads to a 7-fold enrichment of RPS6 mRNA, a 4.7-fold enrichment of RPL32 mRNA, a 11.7-fold enrichment of LDHA mRNA and a 13-fold enrichment of β-actin mRNA (**Fig. 4C**). Similar fold changes are observed when LARP1 immunoprecipitation data is normalized to steady-state input mRNA levels (**Fig. 4E**). As observed for rapamycin, torin1 also significantly (p-value <0.001) enhances the association of both TOP and non-TOP mRNAs to LARP1 (**Fig. 4C** and **Fig. 4E**). Remarkably, torin1 appears to enhance mRNA association to LARP1 more efficiently than rapamycin; this is particularly evident when LARP1 IP data is normalized to steady-state mRNA input levels: for example, rapamycin induces an 8.5-fold enrichment of RPS6 mRNA, while torin1 leads to a 20.1-fold enrichment of the same transcript in LARP1 IPs (**Fig. 4E**). A similar pattern was observed for the remaining mRNAs tested. It appears, therefore, as though torin1 is more effective than rapamycin at enhancing binding to mRNA. The allosteric AKT inhibitor (MK2206) and the non-specific PI3K inhibitor (LY294002) also enhanced the binding of TOP and non-TOP mRNAs to LARP1 protein, albeit less efficiently than rapamycin or torin1 (**Fig. 4C** and **Fig. 4E**). This is consistent with the notion that PI3K/AKT function upstream of mTORC1.

**FIGURE 4.**
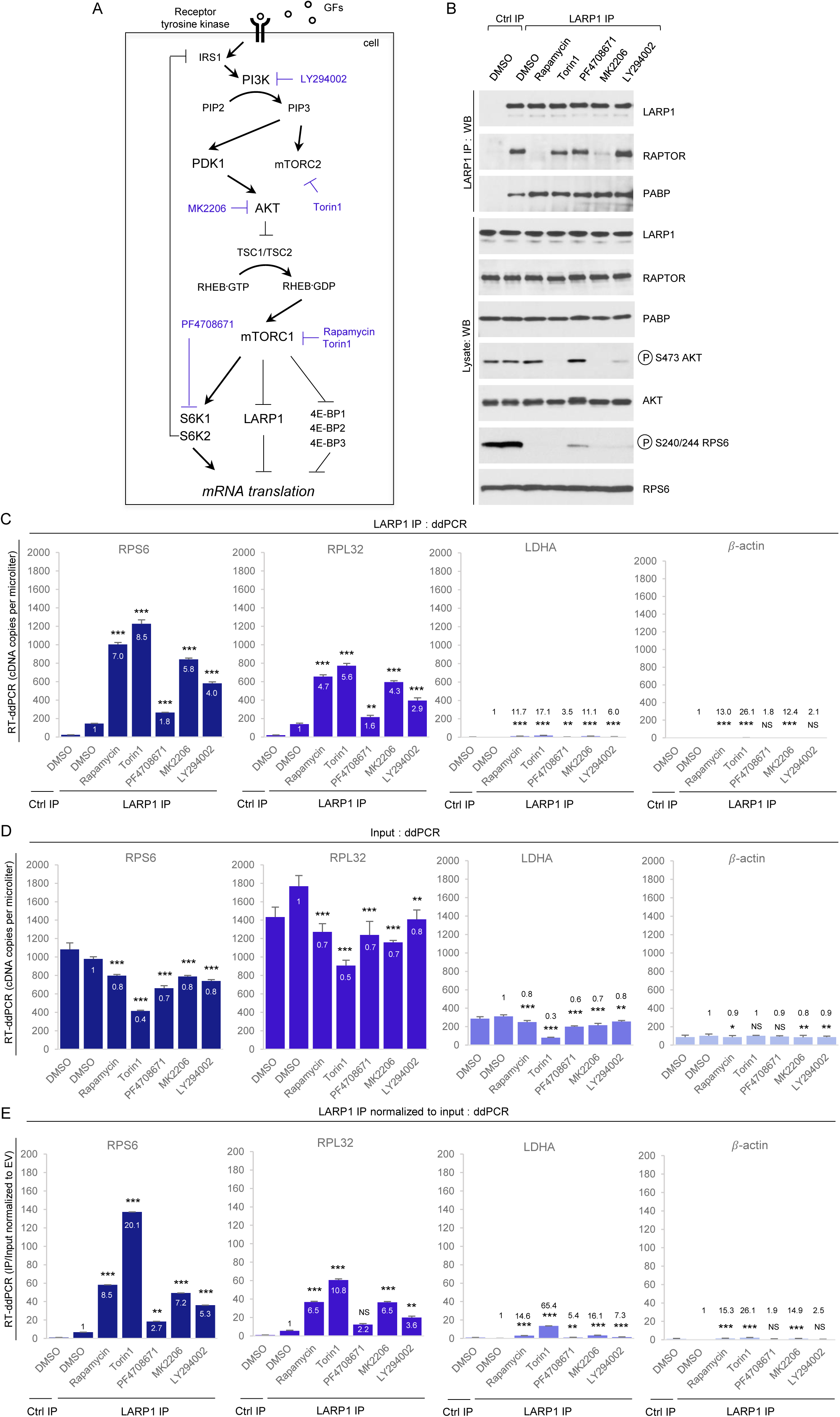
Chemical inhibition of the PI3K/AKT/mTORC1 pathway enhances the binding of endogenous LARP1 to TOP and non-TOP mRNAs. (A) Diagram depicting the PI3K/AKT/mTORC1 signaling pathway and pharmacological agents targeting specific kinases within this pathway. (B-E) HEK293T cells were stimulated with complete media for 3 h in the presence of DMSO (vehicle) or the indicated drugs (see *Experimental Procedures* section for concentration details). Cells were lysed in CHAPS-based extraction buffer in the absence of RNase A and subjected to protein analysis by SDS-PAGE/Western blot (B) or RNA analysis by RT-ddPCR (C-E). *Numbers* within or above bars denote fold enrichment over DMSO/LARP1 IP sample. Asterisks reflect statistical significance: * denotes *p-value* <0.05, ** denotes *p-value* <0.01 and *** denotes *p-value* <0.001. NS signifies non-significant.

It was previously shown that LARP1 associates with mTORC1 (19), and while rapamycin and torin1 both regulate (enhance) LARP1’s association with TOP and non-TOP mRNAs it is unclear whether mTORC1 does so directly or indirectly through a downstream kinase. Ribosomal protein S6 kinases (S6K1 and S6K2) lie downstream of mTORC1 and S6K1 have been previously shown phosphorylate LARP1 (at least *in vitro*) (50). To determine whether S6K1 plays a role in the binding of LARP1 protein to TOP and non-TOP mRNAs, we monitored the effects of PF4708671, a specific S6K1 inhibitor (68), on the binding of endogenous LARP1 to TOP and non-TOP mRNAs. In contrast to rapamycin and torin1, PF4708671 had only a minor effect on LARP1 binding to TOP and non-TOP mRNAs (**Fig. 4C** and **Fig. 4E**). PF4708671 treatment leads to rather small (1.8-fold, 1.6-fold, 3.5-fold and 1.8-fold), and in some instances statistically insignificant, increases in LARP1 binding to RPS6, RPL32, LDHA and β-actin mRNAs, respectively (**Fig. 4C** and **Fig. 4E**). This observation is consistent with the fact that S6Ks have been previously established *<u>not to</u>* play a role in TOP mRNA translation (6,8,12,71).

Since rapamycin (but not torin1) impairs the binding of LARP1 to mTORC1 (**Fig. 1E**), in parallel to the RNA LARP1 IPs, we also performed protein LARP1 IPs and monitored the effect of aforementioned drugs on the interactions between endogenous LARP1 and RAPTOR. As previously observed in **Fig. 1E**, treating cells with rapamycin disrupted the interaction between LARP1 and RAPTOR while torin1 had only a minor (if negligible) effect on the interaction between these proteins (**Fig. 4B**). Neither the S6K1 inhibitor (PF4708671) nor the low-specificity PI3K inhibitor (LY294002) reduce the association of LARP1 with RAPTOR (**Fig. 4B**). By contrast, the allosteric AKT1 inhibitor (MK2206) decreased the interaction between LARP1 and RAPTOR but not that of LARP1 with PABP. Given that rapamycin, torin1, MK2206 and LY294002 all increase the binding of LARP1 to TOP mRNAs to comparable extents (Fig. 4C-E) but only two of these drugs (rapamycin and MK2206) visibly impair LARP1-RAPTOR binding, suggesting that dissociation of LARP1 from RAPTOR is not required for it to bind TOP mRNAs.

### LARP1 is phosphorylated in a rapamycin- and torinl-dependent manner

Having established that the PI3K/AKT/mTORC1 pathway plays an important role in regulating the binding of LARP1 to mRNA, we delved further into the precise molecular mechanism by which this occurs. Using a number of complementary biochemical assays, next we investigated whether the PI3K/AKT/mTORC1 pathway phosphorylates LARP1. First, we used isoelectric focusing (IEF) coupled to sodium-dodecyl sulfate polyacrylamide gel electrophoresis (SDS-PAGE) and Western blot to investigate the effects of rapamycin and torin1 on the electric charge of endogenous LARP1 protein. To this end, HEK293T cells were stimulated with full growth medium in the presence of 0.1 % (v/v) DMSO (vehicle), 100 nM rapamycin or 300 nM torin1 for 3 h and then lysed as described in detail in the *experimental procedures* section. Proteins (from lysates) were fractionated according to their isoelectric point on a ReadyStrip IPG strip (pH range 3–10) and subsequently resolved on a denaturing SDS-PAGE and probed for endogenous LARP1 by Western blot with a LARP1-specific antibody (**Fig. 5A**). LARP1 isolated from cells treated with rapamycin or torin1 showed altered mobility along the IPG strip; specifically, treatment with these drugs raises the isoelectric point (pI) of LARP1 *i. e*. the protein reaches a net charge of zero at higher pH (**Fig. 5A**). The increase in pI denotes a reduction in negative charge of LARP1, possibly reflecting a decrease in phosphorylation. To confirm that the changes in electric charge reflect changes in phosphorylation, we monitored the effect of rapamycin and torin1 on the phosphorylation of LARP1 by *in vivo* orthophosphate metabolic labelling (**Fig. 5B**). Treatment with rapamycin and torin1 (more potently so) decreased the phosphorylation of endogenous LARP1 (**Fig. 5B**) confirming that, indeed, mTORC1 activation status influences LARP1 phosphorylation status. To confirm that rapamycin and torin1 efficiently impaired mTORC1 activity in this particular experiment, we probed lysates with phospho-specific antibodies against S6K1 (phosphorylated on T389) and 4E-BP1 (phosphorylated on T37/46) (**Fig. 5C**). As expected, rapamycin obliterated phosphorylation of T389 on S6K1 and partially reduced the phosphorylation of T37/46 on 4E-BP1, while torin1 abolished the phosphorylation of both substrates.

**FIGURE 5.**
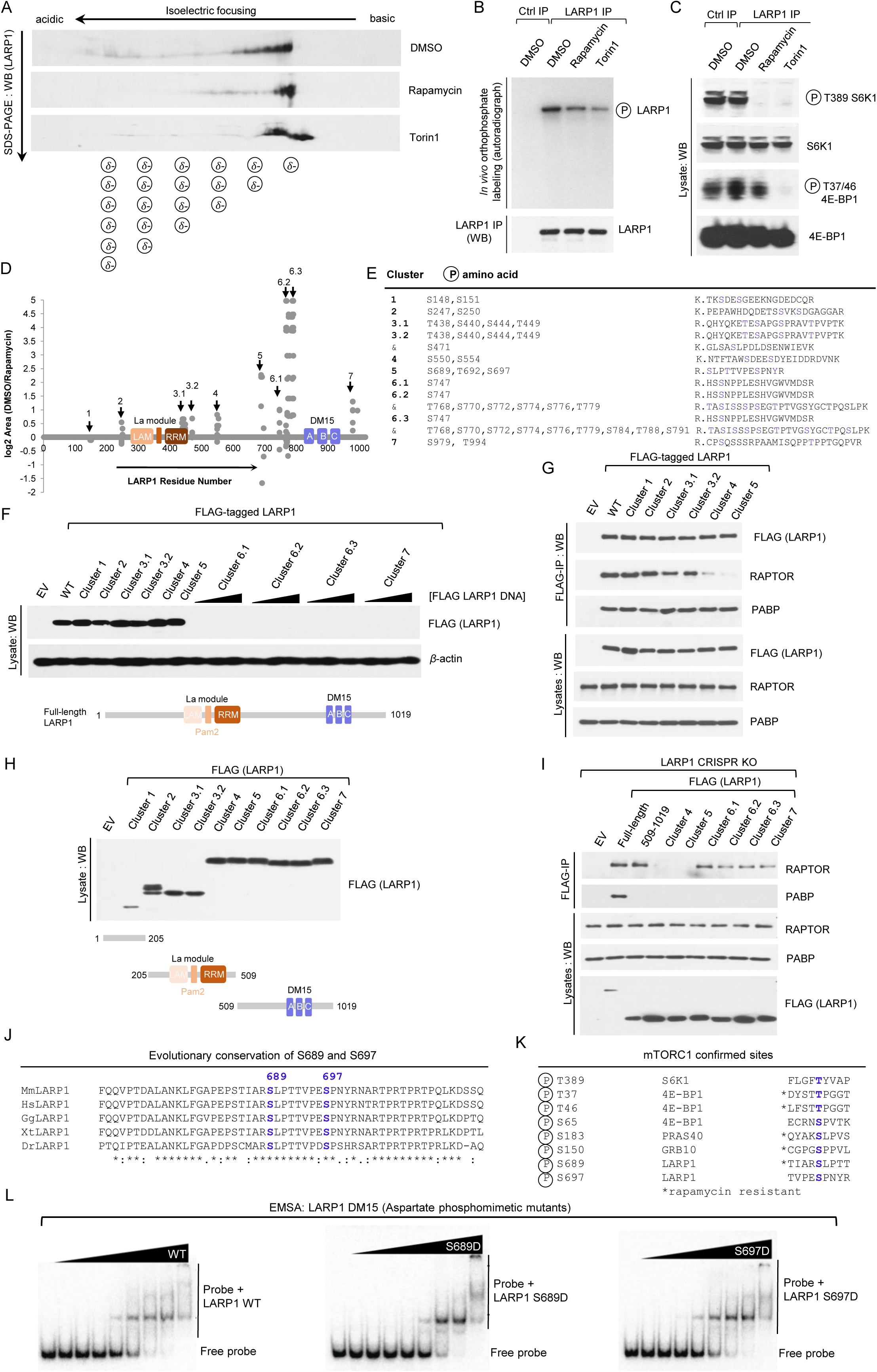
Phosphorylation of S689 regulates LARP1 binding to the 5’UTR of RPS6. (A) Isoelectric focusing of endogenous human LARP1 from HEK293T cells stimulated with complete media for 3 h in the presence of 0.1 % (v/v) DMSO (vehicle), 100 nM rapamycin or 300 nM Torin1. (B,C) Orthophosphate labeling of HEK293T cells stimulated with complete media for 3 h in the presence of 0.1% (v/v) DMSO (vehicle), 100 nM rapamycin or 300 nM Torin1. Cells were lysed and endogenous LARP1 immunoprecipitated with anti-LARP1-specific antibody as described in the *Experimental Procedures* section. Immunoprecipitates and lysates were analysed by SDS-PAGE/Western blot and autoradiography. (D) Graphical representation of rapamycin-sensitive human LARP1 phosphorylated peptides. *X axis* denotes residue number of human LARP1 (isoform 1019aa). *Y axis* denotes log2 area (DMSO/rapamycin). (E) Human LARP1 (1019aa isoform) phosphorylation sites and corresponding tryptic peptides. (F) HEK293T cells were transiently transfected with wildtype FLAGtagged LARP1 or phospho-alanine mutants, lysed and lysates analyzed by SDS-PAGE/western blot. (G) HEK293T cells were transiently transfected empty vector (EV) plasmid DNA or plasmid DNA encoding full-length wildtype human LARP1 or phosphomutants of LARP1 (in which every phosphoresidue in clusters 1, 2, 3.1, 3.2, 4 or 5 was mutated to alanine). Cells were then stimulated with serum for 3 h and subsequently lysed in CHAPS-based buffer and subjected to immunoprecipitation with anti-FLAG antibody. Lysates and FLAG-IP were analyzed by SDS-PAGE/western blot with the indicated antibodies. (H) HEK293T cells were transiently transfected with empty vector (EV) or FLAG-tagged fragments of LARP1 spanning residues 1–205 aa, 205–509 aa or 509–1019 aa, lysed and lysates analyzed as described in (F). (I) HEK293T cells deleted for LARP1 by CRISPR/CAs9 (Clone #5) were transiently transfected with the indicated constructs, stimulated with serum for 3h, lysed in CHAPS based buffer an lysates subjected to immunoprecipitation with anti-FLAG antibody. IP and lysates were analysed by SDS-PAGE/western blot with the indicated antibodies. (J) Alignment of LARP1 protein sequences from Mus musculus (Mm), Homo sapiens (Hs), Gallus gallus (Gg), Xenopus tropicalis (Xt), Danio rerio (Dr) flanking S689 and S697. (K) Confirmed mTORC1 phosphorylation sites on various substrates and degree of rapamycinsensitivity. (L) RPS6 RNA-EMSAs for wildtype DM15, DM15 (S689D) and DM15 (S697D).

### Rapamycin modulates the phosphorylation of 26 serine and threonine and 3 tyrosine residues broadly distributed into 7 clusters

Having established that mTORC1 inhibition (by rapamycin) decreases the phosphorylation of LARP1, we set out to comprehensively identify which are the rapamycin-sensitive residues by liquid chromatography-tandem mass spectrometry (LC-MS/MS). These efforts culminated with the identification of 26 serine, threonine, and 3 tyrosine residues (**Fig. 5D** and **Fig. 5E**); Interestingly, from both functional- and evolutionary-standpoints, most of the rapamycin-sensitive phospho-serine and -threonine residues identified in our screen are conserved across higher eukaryotes – from *D. rerio* to *H. sapiens* (**Supplemental Fig. 2**). This is in contrast with the 3 rapamycin-sensitive phospho-tyrosines identified, which are not conserved in *D. rerio* (**Supplemental Fig. 2**). Although there is precedent for tyrosine phosphorylation (72,73), mTOR is primarily regarded as a serine/threonine kinase. For these reasons, we focused our efforts on understanding the role of rapamycin-sensitive phosphorylation of serine and threonine residues on LARP1. At first glance, we noted that, in some instances, rapamycin impaired the phosphorylation of most of the rapamycin-sensitive residues identified, while in other cases it appeared to enhance phosphorylation of other residues (**Fig. 5D**). Compare, for example, phosphorylated residues within cluster 5 – for some of which its phosphorylation is diminished (above *x* axis) and, in other cases, enhanced by rapamycin treatment (below *x* axis) (**Fig. 5D**). Moreover, rapamycin affected the phosphorylation to different extents (**Fig. 5D**). Prominently, the phosphorylation of residues in cluster 6 (further subdivided into cluster 6.1, 6.2 and 6.3) is exquisitely sensitive to the inhibitory effects of rapamycin, as denoted by the amplitude of the log2 area (DMSO/rapamycin) (**Fig. 5D**), while rapamycin has minor inhibitory effects on clusters 2 and 3 (further subdivided into clusters 2.1 and 2.2). We sought to understand the biochemical significance of LARP1 phosphorylation. The interaction of LARP1 with the mTORC1 is disrupted by rapamycin (**Fig. 1E** and (19)). Similarly, depriving cells of amino acids also reduces the association of LARP1 with mTORC1 (19) – albeit not to the extent that rapamycin does. This raised the intriguing possibility that mTORC1 signaling-mediated phosphorylation of LARP1 regulates the interaction between LARP1 and mTORC1. To test this, we generated alanine-mutant versions of each rapamycin-sensitive phosphorylation site for each cluster individually and tested their expression in HEK293T cells. While mutation of most phosphorylated residues to alanine did not interfere with the expression of LARP1, mutation of clusters 6 or 7 (flanking the DM15 motif) to alanine abrogated the steady-state expression of LARP1 (**Fig. 5F**). These clusters could not, therefore, be tested in subsequent experiments. We tested the effect of replacing serine and threonine residues on the interaction between transiently expressed FLAG-tagged LARP1 and endogenous RAPTOR or PABP. Mutation of clusters 4 (comprising phospho-residues S550 and S554) and, especially, cluster 5 (comprising residues S689, T692 and S697) compromised the binding of ectopic LARP1 to endogenous RAPTOR but not its binding to PABP (**Fig. 5G**). We wished to validate these results and overcome our inability to test the role of the C-terminal phosphorylation sites (clusters 6 and 7) on RAPTOR binding. To address this, we generated fragments of LARP1 spanning the N-terminal region (amino acids 1–205), the mid-domain (amino acids 205–509) and the C-terminal region (amino acids 509–1019) bearing alanines in place of the identified rapamycin-sensitive phosphorylated-serines and – threonines (**Fig. 5H**). Encouragingly, these constructs were successfully transiently overexpressed in HEK293T cells – thus, overcoming the limitation of using full-length LARP1 constructs (*cf*. **Fig. 5F** and 5H). Spurred on by this result, next we tested the interaction between full-length wildtype LARP1 against that of a wildtype fragment of LARP1 spanning amino acids 509–1019 and equivalent fragments of LARP1 bearing alanine mutations in clusters 4, 5, 6.1, 6.2, 6.3 and 7 and endogenous RAPTOR and PABP. To test this, we transiently overexpressed the aforementioned constructs in HEK293T cells deleted for *LARP1* by CRISPR/Cas9 (**Fig. 5I**). CRISPR/Cas9 *LARP1* gene knockout (KO) HEK293T clonal cell lines were generated as shown in **Supplemental Fig. 5A** to 5D and detailed in the *Experimental Procedures* section. CRISPR/Cas9 *LARP1* KO cells were elected as the cell line of choice to test the interaction between phosphorylation-defective fragment mutants of LARP1 to avoid competition between said constructs and endogenous LARP1 for RAPTOR binding. This is particularly important in light of our earlier observation that wildtype fragment 509–1019 binds RAPTOR more weakly than full-length wildtype LARP1 (**Fig. 1D**). Consistent with the results obtained in **Fig. 5G**, substitution of the rapamycin-sensitive phosphorylated-serine and -threonine residues in clusters 4 and 5 by a non-phosphorylatable residue (in this particular case, alanine) impaired the binding of LARP1 to RAPTOR (**Fig. 5I**). By contrast, substitution of the rapamycin-sensitive phospho-serines and -threonines in clusters 6.1, 6.2, 6.3 and 7 had only minor effects on the interaction between LARP1 and RAPTOR. Collectively, these data suggest that phosphorylation of key serine/threonine residues within clusters 4 and 5 (and not in clusters 1, 2, 3.1, 3.2, 6.1, 6.2, 6.3 and 7) regulate LARP1 binding to RAPTOR.

As alluded to earlier, the majority of the rapamycin-sensitive serine and threonine phospho-residues identified by mass spectrometry are conserved in evolution (all the way from *Danio rerio* to *Homo sapiens*) – in some instances, serine to threonine (or *vice-versa*) substitutions can be observed across metazoa suggesting there was evolutionary pressure to maintain phosphorylation of those residues (**Supplemental Fig. 2**). Without exception S689, T692 and S697 (the residues that are most important for the interaction with RAPTOR within cluster 5) are also conserved in evolution (**Fig. 5J**). S689 and S697 are particularly interesting because their surrounding sequences resemble those of other established mTORC1 targets (**Fig. 5K**). The peptide sequence flanking S689 in LARP1 resembles that of S183 in PRAS40 (44,45), a rapamycin-sensitive direct mTORC1 phosphorylation site; while, S697 is followed by a proline at position P698, similar to other proline-directed mTORC1 sites in 4E-BP1 (**Fig. 5K**) (34,35). Having observed that rapamycin inhibits the phosphorylation of S689 and S687 (**Fig. 5D**; see also spectral data in **Supplemental Fig. 2B**), while concurrently enhancing the binding of LARP1 to TOP mRNAs (**Fig. 4C** and **Fig. 4E**), we asked whether these particular phosphoresidues could also play a role in regulating the binding of LARP1 to TOP mRNAs. To answer this question, we bacterially-expressed a fragment of human LARP1 comprising an extended version of the DM15 motif (21,22) spanning residues (665–947) that encompassed S689 and S697 or versions that contained an aspartate in place of either of these residues (an aspartate residue was chosen as phospho-mimetic because it more closely resembles serine than glutamate). Each of the purified fragments was then incubated at various concentrations with a radiolabeled RNA spanning the entire 5’UTR of RPS6 mRNA (capped and methylated at its 5’end) and binding affinity was determined by electrophoretic mobility shift assay (**Fig. 5L**), as previously described in (22). As shown in **Fig. 5L**, wildtype LARP1 (dephospho) fragment efficiently bound to m^7^G-RPS6-UTR radiolabeled probe, as did the S697D mutant; Mutation of S689 to an aspartate reduced the affinity of LARP1 to said probe, as noted by the retention of free probe at higher protein concentrations (**Fig. 5L**). These data suggest that mTORC1-mediated phosphorylation of LARP1 at S689 (but not that of S697) reduces the binding of the extended DM15 LARP1 fragment to the m^7^Gppp cap, the 5’TOP motif, and the 5’UTR of RPS6 mRNA. These data demonstrate that mTORC1 signaling-mediated phosphorylation of LARP1 regulates the RNA binding activity of the latter.

## Discussion

mTORC1-LARP1 is a novel signaling axis with an important role in the control of TOP mRNA translation reviewed in Fonseca et al. (24). LARP1 was originally proposed to enact its inhibitory effects on TOP mRNA translation by competing with eIF4G1 for access to the 5’end of TOP mRNAs (19). Compelling new biochemical and structural studies (22,23) support this model and extend on the preliminary model. Structural data show that LARP1 binds the m^7^Gppp cap moiety and the adjacent 5’TOP motif (21,22); in doing so, LARP1 blocks the loading of the eIF4F complex (comprising eIF4E, eIF4G1 and eIF4A) onto the 5’UTR of TOP transcripts (22), thereby inhibiting cap-dependent translation of TOP mRNAs (23). mTORC1 has long been appreciated to regulate TOP mRNA translation (2,6) but the effector that mediated this effect remained unidentified until recently (19). LARP1 binds to mTORC1 and its depletion renders cells partially resistant to the inhibitory effects of rapamycin and torin1 on TOP mRNA translation (19,23). In this study, we advance our understanding of the mechanism by which mTORC1 controls LARP1 mRNA-binding activity.

LARP1 is phosphorylated *in vivo* in a rapamycin-dependent manner at multiple residues, first documented in two proteome-wide phosphoscreens aimed at identifying novel signaling pathways downstream of mTORC1 (47,48). A follow up study by Kang et al. (49) – aimed at understanding mTORC1 substrate-specificity and rapamycin-sensitivity – revealed that mTORC1 can directly catalyze the phosphorylation of two residues on LARP1 *in vitro:* S766 and S774 (amino acid numbering according 1096 amino acid-long isoform – the corresponding numbering on the 1019 aa-long isoform used in our study and hereafter for simplicity are S689 and S697). S689 and S697 are both equally sensitive to torin1 but exhibited differential sensitivity to rapamycin: S689 is reportedly largely resistant while S697 is sensitive to this drug (49). This conclusion is somewhat surprising because the sequence flanking S689 resembles that of S183 on PRAS40 (a rapamycin sensitive site (44,45)) while S697 is a proline-directed site identical to those found in 4E-BP1 – whose phosphorylation is largely rapamycin resistant (74). A recent study by Hong et al. (50) delved deeper into the LARP1 phosphorylation by mTORC1; their study confirmed that LARP1 is a direct substrate of mTOR *in vitro*. Specifically, using a GST-LARP1 fragment spanning residues 654–731 bearing a double mutation on S689A/T692A they show that mTOR-mediated phosphorylation is lost. These data suggest that S689 and/or T692 are the primary mTOR phosphorylation sites on LARP1 (50).

LARP1 belongs to the LARP superfamily of proteins that comprises 7 members: LARP1, LARP2 (also known as LARP1B), La (sometimes referred to as LARP3), LARP4, LARP5 (also known as LARP4B), LARP6 and LARP7. LARP2 is the closest homolog of LARP1 (with 60% amino acid sequence identity and 73% amino acid sequence similarity). Despite their high degree of sequence conservation, here we show that while both LARP1 and LARP2 bind PABP (through a conserved La module) only LARP1 is capable of strongly interacting with mTORC1. In our hands, myc/Flag-LARP2 <u>does not</u> interact with mTORC1. A recent report by Hong and colleagues (50) proposed that myc-LARP2 interacted weakly with Flag-mTOR/HA-RAPTOR; however, this may be due to simultaneous overexpression of mTOR, RAPTOR and LARP2. Although LARP2 does not interact with endogenous mTORC1, many of the serine/threonine rapamycin-sensitive phosphorylation sites on LARP1 are also conserved in LARP2 (*e.g*. every serine/threonine phospho-residue in clusters 2 through 5 is conserved in LARP2, **Supplemental Fig. 3**) thus raising the intriguing possibility that LARP2 is also a phosphoprotein. Similarly, all-but-one-residue (T779) in cluster 6 of LARP1 are also conserved in LARP2; the latter protein bears an alanine in place of said threonine. Critically, some of the conserved LARP1/LARP2 phospho-residues show conservative serine to threonine or threonine to serine substitutions between LARP1 and LARP2; for instance, T449 in cluster 3 of LARP1, which is a serine in LARP2. Conservative substitutions suggest that, despite sequence divergence between these two closely-related proteins, there was evolutionary pressure to retain phosphorylation. The identity of the kinase(s) that phosphorylates LARP2 is unknown at present, as is the physiological significance of LARP2 phosphorylation. Does LARP2 also bind TOP mRNAs and, if so, does it regulate their translation or stability (as is the case for LARP1 (19,75))? Considerable additional work is required to elucidate these and other questions.

Lastly, in the present manuscript we show that LARP1 employs distinct regions to bind PABP and mTORC1 (see Graphical Abstract for schematic representation). LARP1 binds to PABP via the mid domain region that comprises the La module and to mTORC1 via the DM15 domain. The observation that LARP1 interacts with PABP via the La module (that comprise the La motif, the Pam2 motif and the RNA-recognition motif) is consistent with our earlier observation that the Pam2 motif – nudged between the LaM and the RRM_L5_ is essential for the interaction of LARP1 with PABP (19). The interaction between LARP1 and mTORC1 is more complex; the C-terminal region spanning residues 509–1019 (that comprises the DM15 region and some adjacent sequences) is important for the interaction with RAPTOR (and by extension mTORC1). These findings are consistent with a recent study by Thoreen and colleagues (23) in which the authors report that amino acids 497–1019 on LARP1 mediate RAPTOR binding. The question remained as to which specific residues in LARP1 are required for binding to RAPTOR. Previously, we had proposed that the FRLDI in LARP1 may function as a putative TOS motif (19,61,76), but our earlier data were inconclusive primarily because mutation of the critical phenylanine within LARP1’s putative TOS motif did not affect its binding to mTORC1 (19). In this study, we extended our analysis on the contact points between LARP1 and mTORC1. Our data confirmed that the FRLDI *<u>is not</u>* a TOS motif. Substitution of F889 for an alanine or replacement of the entire motif for the LARP2 (FRREI) sequence did not interfere with the binding of LARP1 to

RAPTOR. Fortuitously, while analyzing this and other putative TOS sequences we observed that mutation of R840 to a glutamate almost abolished RAPTOR binding. We have previously shown that R840 residue plays a critical role in guiding the 5’TOP sequence through a positively charged channel within the DM15 domain (22). Critically this residue also appears to be necessary, but likely not sufficient, for LARP1’s interaction with mTORC1, based on the observation that R840 is conserved in LARP2, yet this protein does not interact with mTORC1. In conclusion, this study delves deeper into the molecular mechanisms by which mTORC1 interacts with and regulates the RNA-binding activity of LARP1. Careful study of this novel signaling node of mTORC1 will advance our understanding of how mTORC1 controls ribosome production. Collectively, these findings may unravel important therapeutic opportunities for the treatment of diseases characterized by dysregulated mTORC1 signaling.

## Experimental Procedures

### Mammalian cell culture, transfection and lysis

HEK293T cells were used in every experiment shown. Cells were cultured/treated in 10-cm tissue culture-treated polystyrene dishes (Corning, catalogue no. 430167) at 37°C in a humidified incubator at 5% CO_2_. Dulbecco’s Modified Eagle’s Media (DMEM) High Glucose (HyClone GE Healthcare, catalogue no. SH30022.01) supplemented with 10% (v/v) fetal bovine serum (Millipore Sigma, catalogue no. F1051) and 100 units/mL penicillin/streptomycin (HyClone GE Healthcare, catalogue no. SV30010) – designated here for ease as *complete media* - was used for cell propagation and treatments. For experiments requiring activation of mTORC1 cells were propagated to near-confluency (~80%) in *complete media*, at which point the media was aspirated and replenished with fresh *complete media* for 3 h. Where indicated cells were simultaneously treated (3 h) with 100 nM rapamycin (LC laboratories, catalogue no. R-5000), 300 nM torin1 (Tocris, catalogue no. 4247), 10 μM PF-4708671 (Tocris, catalogue no. 4032), 10 μM MK-2206 (Cayman Chemicals, catalogue no. 11593) or 30 μM LY294002 (LC laboratories, catalogue #L-7962) or 0.1% (v/v) dimethyl sulfoxide (DMSO) (Millipore Sigma, catalogue no. D1435). DMSO was used as the solvent in the resuspension of every chemical listed above. Where indicated cells were transiently transfected with plasmid DNA for mammalian expression using lipofectamine 2000 reagent (Invitrogen by Thermo Fisher Scientific, catalogue no. 11668-019) as per manufacturer’s instruction. Typically, 4 to 8 μg of plasmid DNA were used to transfect a 10-cm petri dish of near-confluent HEK293T cells. Cells were transfected by incubating the plasmid DNA/lipofectamine 2000 mix in Opti-MEM I (Invitrogen by Thermo Fischer Scientific, catalogue no. 22600-050) for 3 to 4h at 37°C and 5% (v/v) CO_2_ humidified incubator. Transfected cells were then incubated in *complete media* for 24 h followed by another media change for 3 h in *complete media* to activate mTORC1. After mTORC1 stimulation by media change, cells were washed in 5 ml sterile ice-cold phosphate buffered saline (PBS) (important to incline the plate and aspirate all the PBS such that it does not dilute out the lysis buffer) and subsequently lysed in 1 ml of extraction buffer (40 mM HEPES (pH 7.5, room temperature), 0.3% (w/v) CHAPS zwitterionic detergent, 120 mM NaCl, 1mM EDTA, 10 mM sodium pyrophosphate, 10 mM β-glycerophosphate, 50 mM sodium fluoride, 1.5 mM sodium orthovanadate, 1 to 100 μg/mL RNAse A^5^ (Millipore Sigma, catalogue no. 10109169001), 1 mM dithiothreitol (DTT) and cOmplete™ Mini EDTA-free protease inhibitor mixture tablets (Millipore Sigma, catalogue no. 04693159001) for 1 h at 4°C. CHAPS detergent is considerably weaker than most other detergents and, as such, cells <u>must</u> be incubated with extraction buffer (for 1h) before scraping for efficient lysis. Cells were scraped and lysates pre-cleared by centrifugation at 16,000 *g* for 10 min at 4°C. Supernatant was collected onto a fresh microfuge tube. SDS-PAGE lysate samples for Western blot analysis were prepared by adding 50 μl of 4x sample buffer to 150 μl of lysate.

### RNA- and protein-immunoprecipitation

HEK293T cell lysates were prepared as described above. One important difference between RNA- and protein-IPs is that RNAse A was omitted from the extraction buffer for RNA immunoprecipitation but not for protein immunoprecipitation. In addition, 750 μl of lysate was used for RNA immunoprecipitation and 500 μl of lysate was used for protein immunoprecipitation. RNA and protein immunoprecipitation were carried out as follows: 5 μl of LARP1 antibody (AbCam, catalog no. 86359) were added to lysates and incubated for 1 h 30 min rotating end-over-end at 4°C. Then added 35 μl of pre-washed protein G-conjugated magnetic Dynabeads (Life Technologies by Thermo Fisher Scientific, catalogue no. 10003D) to the antibody/lysate mixture and incubated for 1 h rotating end-over-end at 4°C. Following the 1 h incubation, the beads were pelleted by centrifugation at 1000 *g* for 5 min on a table top centrifuge at 4°C, the supernatant aspirated and collected for analysis of unbound material. The beads were then washed twice with 1 ml of extraction buffer. After washing, 500 μl of Trizol reagent (Life Technologies by Thermo Fisher Scientific, catalog no. 15596018) followed by vortexing for 5 secs. Then added an identical volume (500 μl) of extraction buffer to the mix, such that the volume of Trizol to aqueous phase is 1:1 (adding Trizol reagent to the beads prior to adding extraction buffer allows for maximal recovery of RNA from beads) and vortexed samples for an additional 5 secs. Samples were stored overnight at −80°C.

### RNA extraction, cDNA synthesis and reverse transcription-digital droplet PCR (RT-ddPCR)

Samples were thawed from −80°C at room temperature, 50 μl of chloroform was added and then vortexed for 15 secs followed by incubation for 15 min at room temperature. Then subjected to centrifugation at 21,000 *g* for 15 min at 4°C. The aqueous phase was collected and an equal volume of 100% isopropanol was added (to precipitate total RNA) and samples vortexed for 15 sec. Samples were incubated at -20°C to enhance precipitation. Samples were thawed from -20°C and centrifuged at 21,000 *g* for 15 min at 4°C. Supernatant was subsequently discarded and pellet washed gently with 1 ml 75% (v/v) ice-cold ethanol. Centrifugation step was repeated (21,000× g for 5 min at 4°C) and supernatant discarded. RNA pellet was air dried overnight at room temperature suspended in 100 μl RNAse-free water (Millipore Sigma, catalogue no. W4502-1L) for inputs and 10 μl for immunoprecipitates. Reverse-transcriptase reaction was carried out using the iScript Select cDNA synthesis kit (BioRad, catalogue no. 170-8897) as per manufacturer’s protocol with modifications. Briefly, 4 μl of 5x Select reaction mix were added to 1 μl iScript reverse transcriptase, 2 μl Oligo(dT)20 and 10 μl RNA supplemented with RNAse-free water (Millipore Sigma, catalogue no. W4502-1L) to a final volume of 20 μl. The reaction mix was incubated at 42°C for 1 h, followed by 85°C for 5 min. cDNA reaction was then diluted 500x in RNAse-free water (Millipore Sigma, catalogue no. W4502-1L) prior to analysis by digital droplet PCR (ddPCR). Each ddPCR reaction was carried by adding 10 μl QX200^TM^ ddPCR EvaGreen Supermix (BioRad, catalogue no. 186-4034), 0.2 μl of each primer at a stock concentration of 10 μM, 8 μl of diluted cDNA and 1.6 μl RNAse-free water (Millipore Sigma, catalogue no. W4502-1L) to a final reaction volume of 20 μl. The reactions mixtures were transferred to DG8^TM^ Cartridges for QX100^TM^/QX200^TM^ Droplet Generator (BioRad, catalogue no. 186-4008) and 70 μl Droplet Generation Oil for EvaGreen (BioRad, catalogue no. 186-4006). Samples were emulsified on the Droplet Generator and subsequently transferred to a 96-well ddPCR plate. The plate was sealed with aluminum foil and the thermal cycling step ran using the following conditions: 95°C for 5 min, 95°C for 30 secs, ramp down 2°C per second till it reaches 62°C, then ramp up to 95°C and repeat this cycle 45 times. Lastly, samples were cooled down to 4°C for 5 min, heat up to 95°C for 5 min and hold at 12°C indefinitely. Samples were analyzed on Biorad QX200^tm^ Droplet Plate Reader.

### Sodium dodecyl sulfate polyacrylamide gel electrophoresis (SDS-PAGE) and Western blot

Total protein levels and phosphorylation were monitored by SDS-PAGE/Western blot. Samples of lysates were resolved on 10% (w/v) acrylamide (Millipore Sigma, catalogue no. A3553-500G) gel (1.5mm thickness) containing 0.1% (w/v) bis N,N’-methylene bisacrylamide (BioRad, catalogue no. 161-0201) at a ratio of 100:1 of acrylamide to bis N,N’-methylene bisacrylamide. Proteins were then transferred onto a 0.2 μm nitrocellulose membrane (BioRad, catalogue no. 1620112) for 1 h 30 min at 100 V constant by wet (immersion) transfer in a modified Towbin 1x transfer buffer (25 mM Tris, 192 mM glycine at pH 8.3) (77) containing low (10% (v/v)) methanol and 0.1% (w/v) sodium dodecyl sulfate (SDS) for easier transfer of larger molecular weight proteins. The membrane was then blocked in 5% (w/v) defatted milk suspended in Tris buffer saline (1xTBS, 150 mM NaCl, 2.7 mM KCl and 24.7 mM Tris base at pH 7.6) containing 0.02% (v/v) Tween20 (TBS-T) for 1 h at room temperature followed by incubation with primary antibodies in 5% bovine serum albumin (BSA heat shock fraction) (Millipore Sigma, catalogue no. A7906) in TBS-T overnight at 4°C on an orbital shaker. Membranes were then washed twice for 5 min each time in TBS-T and incubated with HRP-conjugated secondary antibodies for 45 min at room temperature on an orbital shaker. Unbound antibody was washed by rinsing membranes thrice in TBS-T for 5 min each time. Protein was detected by enhanced chemiluminescence (ECL) using the Western Lightning Plus-ECL reagent (Perkin Elmer Inc., catalogue no. NEL105001EA). ECL signal was detected by autoradiography using HyBlot CL film (Denville Scientific Inc., catalogue no. E3018). All proteins were analyzed as aforementioned with exception to 4E-BPs; the latter were resolved on 13.5% (w/v) acrylamide gels (1.5mm thickness) containing 0.36% (w/v) bis N,N’-methylene bisacrylamide (BioRad, catalogue no. 161-0201) at a ratio of 37.5:1 of acrylamide to bis N,N’-methylene bisacrylamide). Proteins were then transferred onto a 0.2 μm nitrocellulose membrane (BioRad, catalogue no. 1620112) for 1h at 100V constant by wet (immersion) transfer in a modified Towbin 1x transfer buffer (25 mM Tris, 192 mM glycine at pH 8.3) (77) containing high (20% (v/v)) methanol and 0.1% (w/v) sodium dodecyl sulfate (SDS). Higher methanol percentage is used to avoid over-transfer of molecular weight proteins (such as 4E-BP1). To enhance the retention of 4E-BP1 protein on the membrane, the proteins were crosslinked to the membrane by incubating the membrane for 30 min with 0.05% (v/v) glutaraldehyde solution (Bio Basic Canada Inc., catalogue no. GC3870) prepared in phosphate buffered saline (1x PBS, 137 mM NaCl, 2.7 mM KCl, 10 mM Na_2_HPO_4_, 2 mM KH_2_PO_4_ at pH7.4) on an orbital shaker. Membrane was rinsed twice with deionized water and subsequently blocked with defatted milk, probed with primary/secondary antibodies and developed as described above.

### Antibodies

Anti-human LARP1 rabbit polyclonal antibody (catalogue no. ab86359) and anti-human PABP rabbit polyclonal antibody (catalogue no. ab21060) were purchased from AbCam. Anti-FLAG M2 mouse monoclonal antibody (catalogue no. F1804) and anti-rabbit horseradish peroxidase (HRP)-conjugated IgG (catalogue no. A0545) were purchased from Millipore Sigma. Anti-human RAPTOR rabbit polyclonal antibody (catalogue no. A300-553A) was purchased from Bethyl Laboratories. Anti-human RPS6 (C-8) mouse monoclonal IgG antibody (catalogue no. sc-74459) and anti-human S6K1 (C-18) rabbit polyclonal IgG antibody were purchased from Santa Cruz Biotechnology. Anti-rabbit eIF4E1 mouse IgG antibody (catalogue no. 610269) was purchased from BD Transduction Laboratories. Anti-human mTOR rabbit monoclonal (7C10) antibody (catalogue no. 2983), anti-phospho S473 human AKT rabbit polyclonal antibody (catalogue no. 9271S), anti-human AKT pan rabbit monoclonal (C67E7) antibody (catalogue no. 4691S), anti-hemagglutinin (HA) rabbit monoclonal (C29F4) antibody (catalogue no. 3724), anti-human eIF4G1 rabbit monoclonal (C45A4) antibody (catalogue no. 2469S), anti-phospho T389 human S6K1 rabbit monoclonal (108D2) antibody (catalogue no. 9234), anti-human S6K1, anti-phospho T37/T46 human 4E-BP1 rabbit monoclonal (236B4) antibody (catalogue no. 2855), anti-phospho S240/S244 human RPS6 rabbit polyclonal antibody (catalogue no. 2215S), anti-human 4E-BP1 rabbit monoclonal (53H11) antibody (catalogue no. 9644S) and anti-mouse horseradish peroxidase (HRP)-conjugated IgG (catalogue no. 7076S) antibody were purchased from Cell Signaling Technology. Primary anitbodies were used at 1:1000 dilution, while secondary antibodies were used at 1:10,000.

### Generation of plasmid DNAs

pCMV6-entry human wildtype LARP1 transcript variant 1 (accession number NM_0153315) myc/FLAG-tagged (originally described in (19)) (catalogue no. RC200935) was purchased from Origene. pCMV6-human wildtype LARP2 transcript variant 1 (accession number NM_018078) (catalogue no. RC213675) and transcript variant 3 (accession no. NM_032239) (catalogue no. RC219586) myc/FLAG-tagged were also purchased from Origene. pCMV2 FLAG-tagged human wildtype La (55), pCMV2 FLAG-tagged human wildtype LARP4 (55), pCMV2 FLAG-tagged human wildtype LARP5 (55), human wildtype LARP6 (55) and human wildtype LARP7 were kindly gifted to us by Dr. Richard J. Maraia (Section on Molecular Biology and Cell Biology, Intramural Research Program in Genomics of Differentiation, Eunice Kennedy Shriver National Institute of Child Health and Human Development, National Institute of Health, Bethesda, MD, USA). pCMV6-entry human LARP1 F889A (FRLDI→ARLDI) and DTOS (DFRLDI) mutants have been previously generated and described in (19). pCMV6-entry human LARP1 R840E/Y883A double mutant has been previously generated and described in (22). pCMV5-HA-tagged human wild type PRAS40 isoform b (also known as AKT1S1 for AKT1 substrate 1, accession no. BC007416) was originally cloned and described in (45). The pCMV5-HA-tagged human PRAS40 F129A (FVMDE→AVMDE) and M131A/D132A (FVMDE→FVAAE) (numbering according to human isoform b) were previously generated by site-directed mutagenesis and described in (45).

### Generation of LARP1 CRISPR/Cas9 Knockout (KO) HEK293T cell lines

To create plasmids for expression of LARP1-specific gRNAs, sense (5’-CACCGAGACACATACCTGCCAATCG-3’) and antisense (5’-AAACCGATTGGCAGGTATGTGTCTC-3’) oligonucleotides were annealed and cloned into Esp3I-digested LentiCRISPRv2, resulting in a vector designated LentiCRISPR-LARP1gRNA. HEK293T cells were maintained in DMEM medium (Gibco/Invitrogen by Thermo Fisher Scientific) supplemented with 10% fetal calf serum (Gibco/Invitrogen by Thermo Fisher Scientific) and 100 units/mL penicillin/streptomycin (Gibco/Invitrogen by Thermo Fisher Scientific, catalogue no. 15140122). All cells were cultured at 37°C in 5% (v/v) CO_2_. One day prior to transfection, HEK293T cells were seeded at a density of 3 x 10^5^ cells/well in 6-well plates. Transfections were carried out using 1 μg LentiCRISPR-LARP1gRNA and Lipofectamine 2000 Transfection Reagent (Invitrogen by Thermo Fisher Scientific, catalogue no. 11668-019) according to manufacturer’s protocol. Two days after transfection, the cells were reseeded at a density of 0.2 cells/well in 96-well plates. After expansion of single cells, genomic DNA was purified using the GenElute Mammalian Genomic DNA Miniprep Kits (Millipore Sigma-Aldrich, catalogue no. G1N10) according to manufacturer’s protocol. LARP1 CRISPR/Cas9 KO was verified by PCR on genomic DNA using the primers 5’ -GGGAAAGGGATCTGCCCAAG-3’ and 5’ -CACCAGCCCCATCACTCTTC-3’ and a Pfu Ultra II DNA polymerase (Agilent Technologies) according to manufacturer’s protocol followed by Sanger sequencing of the resulting PCR-product (GATC Biotech) using the primer 5’ -GGGAAAGGGATCTGCCCAAG-3’.

### Site-directed mutagenesis and oligonucleotides

Phosphorylation and RNA-binding mutants of human LARP1 were generated by site-directed mutagenesis using Pfu Ultra HF DNA polymerase (Agilent Technologies, catalogue no. 600380-51) as described in the manufacturer’s protocol. Where applicable, oligonucleotides were designed using the PrimerGenesis software tool for automated oligonucleotide design (www.primergenesis.org). The following oligonucleotides were employed for site-directed mutagenesis of human LARP1 and human LARP2 genes:

LARP1_L891R/D892E_forward (5’-

GGCCTGGAAAAGAAGTTCCGGCGGGAAATATTCAAGGATTTTCAG-3’);

LARP1_L891R/D892E_reverse (5’-

CTGAAAATCCTTGAATATCTCCCGCCGGAACTTCTTTTCCAGGCC-3’);

LARP1_L928A/D929A_forward

(5’CCAAAGCCAAAAATGCGGCCATTGACCCCAAACTGCAAGAATACCTC-3’);

LARP1_L928A/D929A_reverse

(5’GCAGTTTGGGGTCAATGGCCGCATTTTTGGCTTTGGAATATTTCAAG-3’);

LARP1_K924Q/A925S/N927T/L928Q/D929S_forward

(5’TCTTGAAATATTCCCAATCCAAAACTCAGAGCATTGACCCCAAACTGCAAGAATA CCTCG-3’);

LARP1_K924Q/A925S/N927T/L928Q/D929S_reverse

(5’GCAGTTTGGGGTCAATGCTCTGAGTTTTGGATTGGGAATATTTCAAGAAGGCCCAGAAC-3’);

LARP2_R790L/E791D_forward

(5’GGACTGGAAAAAAAATTCAGGCTAGATATTTTTCAGGATTTCCAAGAAGAAACC-3’)

LARP2_R790L/E791D_reverse

(5’CTTCTTGGAAATCCTGAAAAATATCTAGCCTGAATTTTTTTTCCAGTCCATAAC-3’)

LARP1_R840E_forward

(5’GGAGATGAACACACTCTTCGAATTCTGGTCCTTCTTCCTCCG-3’);

LARP1_R840E_reverse

(5’CGGAGGAAGAAGGACCAGAATTCGAAGAGTGTGTTCATCTCC-3’);

LARP1_Y883A_forward

(5’CTACAGTGCTGGCCTGGAAAAGAAGTTCCGGCTGGACATATTC-3’);

LARP1_Y883A_reverse

(5’CCAGGCCAGCACTGTAGTATCTATCGAAAAAGGCACTCCAAACC-3’);

LARP1_S148A/S151A_forward

(5’AAGGAGAGTCCAAAAACCAAAGCAGATGAAGCAGGGGAGGAAAAGAATGGAGATGAGGAT-3’);

LARP1_S148A/S151A_reverse

(5’ATCCTCATCTCCATTCTTTTCCTCCCCTGCTTCATCTGCTTTGGTTTTTGGACTCTCCTT-3’);

LARP1_S247A/S250A_forward

(5’GACCAGGATGAGACATCGGCTGTGAAGGCTGATGGGGCTGGTGGGGCGCGGGCTTCCTTC-3’);

LARP1_S247A/S250A_reverse

(5’GAAGGAAGCCCGCGCCCCACCAGCCCCATCAGCCTTCACAGCCGATGTCTCATCCTGGTC-3’);

LARP1_T438A/S440A/S444/T449A_forward

(5’GTTCCCCGTCAGCACTACCAAAAGGAGGCAGAGGCGGCACCTGGCGCTCCTCGTGCAGTCGCCCCAGTGCCAACCAAAACAGAGGAGGTC-3’);

LARP1_T438A/S440A/S444/T449A_reverse

(5’GACCTCCTCTGTTTTGGTTGGCACTGGGGCGACTGCACGAGGAGCGCCAGGTGCCGCCTCTGCCTCCTTTTGGTAGTGCTGACGGGGAAC-3’);

LARP1_S471A_forward (5’AAGGGCCTGTCTGCCGCCCTGCCTGACCTGGAT-3’);

LARP1_S471A_reverse (5’ATCCAGGTCAGGCAGGGCGGCAGACAGGCCCTT-3’);

LARP1_S550A/S554A_forward

(5’ACCTTCACTGCCTGGGCTGATGAGGAAGCTGACTATGAGATTGAT-3’);

LARP1_S550A/S554A_reverse

(5’ATCAATCTCATAGTCAGCTTCCTCATCAGCCCAGGCAGTGAAGGT-3’);

LARP1_S689A/T692A/S697A_forward

(5’CCCTCCACCATCGCCCGCGCTCTACCAGCCACTGTCCCAGAGGCACCAAACTACCGCGGC-3’);

LARP1_S689A/T692A/S697A_reverse

(5’GCCGCGGTAGTTTGGTGCCTCTGGGACAGTGGCTGGTAGAGCGCGGGCGATGGTGGAGGG-3’);

LARP1_S747A_forward (5’AAGACAAGACACAGTGCAAACCCACCCTTGGAG-3’);

LARP1_S747A_reverse (5’CTCCAAGGGTGGGTTTGCACTGTGTCTTGTCTT-3’);

LARP1_T768A/S770A/S772A/S774A/S776A/T779A_forward

(5’ATGGATTCCCGTGAGCACAGGCCCCGTGCTGCTGCCATCGCCTCCGCCCCCGCAGAAGGGGCGCCTACAGTTGGCAGCTATGGCTGTACC-3’);

LARP1_T768A/S770A/S772A/S774A/S776A/T779A_reverse

(5’GGTACAGCCATAGCTGCCAACTGTAGGCGCCCCTTCTGCGGGGGCGGAGGCGATGGCAGCAGCACGGGGCCTGTGCTCACGGGAATCCAT-3’);

LARP1_S784A/T788A/S791A_forward

(5’GGGACGCCTACAGTTGGCGCCTATGGCTGTGCCCCTCAGGCATTGCCCAAGTTCCAGCAT-3’);

LARP1_S784A/T788A/S791A_reverse

(5’ATGCTGGAACTTGGGCAATGCCTGAGGGGCACAGCCATAGGCGCCAACTGTAGGCGTCCC-3’);

LARP1_S979A_forward (5’AGGAAGCGGTGCCCCGCCCAGTCTTCCAGCAGG-3’);

LARP1_S979A_reverse (5’CCTGCTGGAAGACTGGGCGGGGCACCGCTTCCT-3’);

LARP1_S689D_forward (5’CCTCCACCATCGCCCGCGATCTACCAACCACTGTCC-3’);

LARP1_S689D_reverse

(5’GGACAGTGGTTGGTAGATCGCGGGCGATGGTGGAGG-3’);

LARP1_S697D_forward

(5’CTACCAACCACTGTCCCAGAGGATCCAAACTACCGCAACACCAGG-3’);

LARP1_S697D_reverse

(5’CCTGGTGTTGCGGTAGTTTGGATCCTCTGGGACAGTGGTTGGTAG-3’).

The following oligonucleotides were used to sequence human LARP1 and human LARP2:

LARP1_sequencing1_forward   (5’ATGCTTTGGAGGGTGCTTTTG-3’);

LARP1_sequencing2_forward   (5’GTTCCTAAACAGCGCAAAGGC-3’);

LARP1_sequencing3_forward   (5’TGCCAGCGAGGCGGGCAGAAG-3’);

LARP1_sequencing4_forward   (5’GACCAGGATGAGACATCGAGTG-3’);

LARP1_sequencing5_forward   (5’GTGGATCAGGAACTGCTCAAAG-3’);

LARP1_sequencing6_forward   (5’GAGGAACCAGAAAAGTGGCCTC-3’);

LARP1_sequencing7_forward   (5’ATTGAAGTGAAGAAGAGGCCTC-3’);

LARP1_sequencing8_forward   (5’AGGGATGTCAACAAGATCCTC-3’);

LARP1_sequencing9_forward   (5’GAGCAGTTTGACACACTGACC-3’);

LARP1_sequencing10_forward   (5’TCACGGTTTTACCCAGTGGTG-3’);

LARP1_sequencing11_forward   (5’GAACTGCTCAAGGAAAATGGC-3’);

LARP1_sequencing12_forward   (5’TACAGTTATGGCCTGGAAAAG-3’);

LARP1_sequencing13_forward   (5’CGACACTCAGTGGTAGCAGGAG-3’);

LARP1_sequencing14_reverse   (5’AGGGAATGGCAATGGCTTCTC-3’);

LARP2_sequencing1_forward   (5’ATGGAGAATTGGCCAACACCAAG-3’);

LARP2_sequencing2_forward   (5’AGATGTCAACCTGAAGCAAATAAAC-3’);

LARP2_sequencing3_forward   (5’GATGGTACAGGTGTACAGGTG-3’);

LARP2_sequencing4_forward   (5’CCTCCACGCAGTGTGCCACCAAC-3’);

LARP2_sequencing5_forward   (5’-TGTTCTTCAGAAGAACCAGAAC-3’);

LARP2_sequencing6_forward   (5’GAAAACAAACACACAGCCATAAAG-3’);

LARP2_sequencing7_forward   (5’ACACCCAAAACACCTCGAACAC-3’);

LARP2_sequencing8_forward   (5’CTTTTGAAGGAAAATGGCTTTAC-3’);

LARP2_sequencing9_forward   (5’AAAAAAGACTACGAATCTGGTCAGC-3’);

LARP2_sequencing10_reverse   (5’TGAGCTGTTCCGGGATCCAGG-3’).

The following oligonucleotides were used for the analysis of human RPS6, RPL32, LDHA and β-actin mRNA levels by RT-ddPCR:

RPS6_forward (5’-CTGGGTGAAGAATGGAAGGGTT-3’);

RPS6_reverse (5’-TGCATCCACAATGCAACCAC-3’);

RPL32_forward (5’-AGCCATCTCCTTCTCGGCAT-3’);

RPL32_reverse (5’-TCAATGCCTCTGGGTTTCCG-3’);

LDHA_forward (5’-AAAGGCTACACATCCTGGGC-3’);

LDHA_reverse (5’-GGTGCACCCGCCTAAGATTC-3);

β-actin_forward (5’-TGATGATATCGCCGCGCTC-3’);

β-actin_reverse (5’ -CATCACGCCCTGGTGCC-3’).

### Isoelectric focusing

Briefly, isoelectric focusing was performed as described in manufacturer’s manual. In detail, HEK293T cells were stimulated with *complete media* (as described above) for 3h in the presence of 0.1 (v/v) DMSO, 100 nM rapamycin or 300 nM torin1. *<u>Lysis:</u>* Cells were then lysed by incubating in 800 μl rehydration buffer for ih at 4°C at which point cells were scraped, lysates pre-cleared by centrifugation at 16,000× g for 10min at 4°C. *<u>Sample cleanup:</u>* Samples were then further cleaned by using the ReadyPrep™ 2-D Cleanup Kit: 200 μl of lysate was transferred into a clean microfuge tube, 600 μl precipitation agent 1 were added to lysate and the microfuge tube vortexed. Vortexed sample was incubated for 15 min on ice. 600 μl of precipitation agent 2 were then added to the mixture of lysate and precipitation agent 1 and the microfuge tube vortexed. Samples were then centrifuged at 21,000 *g* for 5 min to form a light pellet. Supernatant was discarded without disturbing the pellet. Residual liquid in the microfuge tube was collected and discarded following centrifugation for 30 secs at 21,000 *g*. 40 μl of wash reagent 1 was added on top of the pellet, tube vortexed and centrifuged 21,000 *g* for 5 min. The wash was then discarded with a pipette and 25 μl of ReadyPrep proteomic grade water (BioRad, catalogue no. 163-2091). Tube was vortexed for 20 secs. 1 ml of wash reagent 2 (pre-chilled at – 20°C) and 5 μl of wash 2 additive was added to the tube. Tube was vortexed for 1 min. Samples in the tube were then incubated at -20°C for 30 min. Samples were vortexed once for 30 secs midway through the incubation. After the incubation period, samples were centrifuged at 21,000 *g* for 5 min to form a tight pellet. The supernatant was discarded, the tube centrifuged briefly for 30 secs to discard any remaining wash. The pellet was air-dried at room temperature for approximately 5 min until it looked translucent. The pellet was then resuspended in 2-D rehydration buffer (see preparation below), vortexed for approximately 3 min (or until the pellet was fully resuspended). Sample was centrifuged at 21,000 *g* for 5min at room temperature to clarify the protein sample and the supernatant used for IEF in IPG strips. *<u>Sample application to IPG strip:</u>* the IPG strip (pH 3-10, 11cm) was thawed from -20°C and the rehydration/sample buffer lyophilized powder reconstituted by adding 6.1 ml of nanopure water supplied with the kit. Apply 185 μl of sample along the back edge of the channel of the rehydration/equilibration tray evenly, leaving 1 cm at each end. This step was repeated for each sample using a different channel. Once the protein samples were loaded into the rehydration tray forceps were used to peel the coversheet from the thawed ReadyStrip IPG strip and the strip was placed over the sample in the rehydration/equilibration tray with the gel side facing down onto the sample. Air bubbles trapped underneath the strip were carefully removed by gently lifting the strip up and down with the forceps or as a last resort gently pressing down the strip. Each strip was then overlayed with 2 ml of mineral oil (BioRad, catalogue no. 163-2129) to prevent evaporation during the rehydration process. Mineral oil was added slowly to the plastic backing of the strips while moving the pipet along the length of the strip to avoid mineral oil seeping beneath the strip. The rehydration/equilibration tray was covered with the plastic lid provided and the tray left sitting on a level bench overnight (11–16 h) at room temperature to rehydrate the IPG strips and load the protein sample. *<u>Isoelectric focusing:</u>* Using forceps, paper wicks were placed at both ends of the Protean IEF focusing tray covering the wire electrodes. 8 μl of nanopure water (provided with kit) was added onto each wick. IPG strips were picked up from rehydration/equilibration tray using forceps and held vertically for 10 secs over filter paper to allow the mineral oil to drain. Draining the oil is important to remove the unabsorbed protein which would otherwise cause horizontal streaking. IPG strips were then transferred to the corresponding channel in the Protean IEF focusing tray with the gel side facing down. Once placed in the focusing tray the strips were again covered with 2 ml of fresh mineral oil. Any trapped air bubbles were removed as described above. The focusing tray was covered with the lid and placed on the Protean IEF cell and the cover closed. Strips were resolved using the following 3-step protocol. Step 1: 250 V for 20 min and linear ramp; Step 2: 8,000 V for 2 h 30 min and linear ramp; Step 3: 8,000 V for an undefined time until it reached 20,000 V-h with rapid ramping. This protocol takes an approximate time of 6 h and accumulated voltage of approximately 30,000 V. A default cell temperature of 20°C with a maximum current of 50 μA/strip and no rehydration parameters were used. *<u>Equilibration of IPG strips:</u>* following completion of the electrophoresis step, the mineral oil from was drained from IPG strips by holding them vertically using forceps for 10 secs over filter paper and the strips were then transferred onto a clean/dry disposable rehydration/equilibration tray with the gel side facing up. Equilibration buffers 1 (containing 6 M urea, 2% (w/v) sodium dodecyl sulfate in 50 mM Tris-HCl buffer pH 8.8 and 10 mg/ml dithiothreitol) was prepared by adding 13.35 mL of 30% (v/v) glycerol solution supplied in kit to buffer. Equilibration buffer 2 (containing containing 6 M urea, 2% (w/v) sodium dodecyl sulfate in 50 mM Tris-HCl buffer pH8.8) was prepared by was prepared by adding 13.35 mL of 30% (v/v) glycerol solution supplied in kit to buffer and 40 mg/ml of iodoacetamide (alkylating agent). Iodoacetamide is added to equilibration buffer 2 to prevent sulfhydryl bond formation between free thiol groups of cysteine residues which interfere with the 2-dimension (2-D) electrophoresis step. Contents were mixed at room temperature using a stir plate until all solids were fully dissolved. 4 mL of equilibration buffer 1 was added to each rehydration/equilibration tray channel containing an IPG strip. Tray was placed on an orbital shaker and gently shaken for 10 min at room temperature. A slow shaker speed was used to prevent the buffer from sloshing out of the tray. At the end of the 10min incubation, equilibration buffer 1 was discarded by tipping the liquid gently from the tray. Once most of the liquid was decanted the tray was flicked to remove the last drops of equilibration buffer 1. Four milliliters of equilibration buffer 2 were then added to each channel of the rehydration/equilibration tray containing an IPG strip and incubated for 10 min at room temperature with shaking (on an orbital shaker). *<u>Second-dimension (2-D):</u>* the second-dimension step (2-D sodium dodecyl sulfate polyacrylamide gel electrophoresis) was performed in 4–15% Criterion™ TGX™ Precast Midi SDS-PAGE gels (11 cm IPG/prep+1 well) (BioRad, catalogue no. 5671081). Briefly, IPG strip was rinsed in a graduated cylinder containing 100 ml of 1xTris/glycine/SDS running buffer. The 4–15% Criterion™ TGX™ Precast Midi SDS-PAGE gel well was rinsed with nanopure water and the excess water blotted using Whatman 3MM paper. The IPG strip was laid gel side onto the back plate of the SDS-PAGE gel. Holding the SDS-PAGE vertically, ReadyPrep Overlay agarose solution (BioRad, catalogue no. 163-2111) was added gently to the well using a Pasteur pipet. Using forceps, the IPG strip was carefully mounted/pushed onto the well, avoiding air bubbles underneath the strip, and the agarose left to solidy for 5 min at room temperature. SDS Proteins were eluted from the strip by adding isoelectric focusing (IEF) gel sample buffer (supplied with kit) containing 50% (v/v) glycerol (BioRad, catalogue no. 161-0763) and coomassie blue R-250 stain for sample visualization. Proteins were resolved by electrophoresis in 1x Tris/glycine/SDS running buffer containing (25 mM Tris, 192 mM glycine, 0.1% (w/v) sodium dodecyl sulfate (SDS) at pH 8.3) at 150 V constant for approximately 1 h 30 min to 2 h. Proteins were then transferred onto a 0.2 μm nitrocellulose membrane (BioRad, catalogue no. 1620112) for 1 h 30 min at 100 V constant by wet (immersion) transfer in a modified Towbin 1x transfer buffer (25 mM Tris, 192 mM glycine at pH 8.3) (77) containing low (10% (v/v)) methanol and 0.1% (w/v) sodium dodecyl sulfate (SDS) for easier transfer of larger molecular weight proteins. Membrane was blocked in 5% (w/v) defatted milk in TBS-T for 1 h at room temperature on an orbital shaker, followed by incubation overnight at 4°C on an orbital shaker with primary anti-human LARP1 antibody (AbCam, catalogue no. 86359) at 1:1000 dilution in 5% (w/v) bovine serum albumin (BSA, heat shock fraction) (Millipore Sigma, catalogue no. A7906) in TBS-T. Unbound primary antibody was washed by incubating membrane twice for 5 min each time in TBS-T. The membrane was subsequently incubated with anti-rabbit horseradish peroxidase (HRP)-conjugated IgG (Millipore Sigma, catalogue no. A0545) for 1 h at room temperature on an orbital shaker, at which point the unbound secondary antibody was washed three times in TBS-T (5 min each time). Protein was detected by enhanced chemiluminescence (ECL) using the Western Lightning Plus-ECL reagent (Perkin Elmer Inc., catalogue no. NEL105001EA). ECL signal was detected by autoradiography using HyBlot CL film (Denville Scientific Inc., catalogue no. E3018).

### Orthophosphate labeling

HEK293T cells were propagated to near-confluency (~80%) in 10 cm tissue culture-treated polystyrene dishes (Corning, catalogue no. 430167) at 37°C in a humidified incubator at 5% (v/v) CO_2_ in phosphate-containing *complete media* (Dulbecco’s Modified Eagle’s Media (DMEM) High Glucose (HyClone GE Healthcare, catalogue no. SH30022.01) supplemented with 10% (v/v) fetal bovine serum (Millipore Sigma, catalogue no. F1051) and 1% (v/v) penicillin/streptomycin (HyClone GE Healthcare, catalogue no. SV30010)). Once cells reached ~80% confluency, the *complete media* was aspirated and replaced with 5 ml of fresh *complete media* containing Phosphorus 32 (^32^P) orthophosphoric acid (Perkin Elmer, NEX053005MC) (~1 mCi were used per 10 cm dish) in the presence of 0.1% (v/v) DMSO (vehicle), 100 nM rapamycin or 300 nM torin1. Cells were incubated with vehicle/drugs for 3h at 37°C in a humidified incubator at 5% (v/v) CO_2_. Washed once in ice-cold phosphate buffered saline (PBS) and lysed in 1 mL radio-immunoprecipitation assay (RIPA) extraction buffer containing 50 mM Tris-HCl (pH 8 at room temperature), 150 mM sodium chloride, 1% (v/v) Igepal CA-630 (Nonidep P40, NP40), 0.5% (w/v) sodium deoxycholate and 0.1% (w/v) sodium dodecyl sulfate (SDS), 50 mM sodium fluoride, 1.5 mM sodium orthovanadate, 1 μg/mL RNAse A (Millipore Sigma, catalogue no. 10109169001), 1 mM dithiothreitol (DTT) and cOmplete^TM^ Mini EDTA-free protease inhibitor mixture tablets (Millipore Sigma, catalogue no. 04693159001). Cells were scraped and lysates pre-cleared by centrifugation at 16,000 *g* for 10 min at 4°C. Supernatants were transferred into a fresh microfuge tube and used for radio-immunoprecitation as follows: 900 μl lysate were incubated with 9 μl of anti-human LARP1 rabbit polyclonal antibody (AbCam, catalogue no. ab86359) for 1 h 30 min rotating end-over-end at 4°C. Then added 35 μl of pre-washed protein G-conjugated magnetic Dynabeads (Life Technologies by Thermo Fisher Scientific, catalogue no. 10003D) to the antibody/lysate mixture and incubate for 1h rotating end-over-end at 4°C. Following the 1h incubation, the beads were pelleted by centrifugation at 1000× g for 5 min on a table top centrifuge at 4°C, the supernatant aspirated and collected for analysis of unbound material. The beads were then washed thrice with 1 ml of RIPA extraction buffer followed by resuspension in 50 μl 4x SDS-PAGE sample buffer and boiling for 5 min at 95°C. Beads were then pelleted and stored at -20°C until further analysis by SDS-PAGE/Western blot/^32^P-autoradiography. 10 μl of immunoprecipitate was used to monitor the phosphorylation of endogenous LARP1 by SDS-PAGE/Western blot/^32^P-autoradiography. SDS-PAGE/Western blot was performed as described above (up until the blocking step). Following blocking the nitrocellulose membrane was enveloped in cling film and exposed for 2 h to 48 h to autoradiography HyBlot CL film (Denville Scientific Inc., catalogue no. E3018) at −80°C and developed using the Konica Minolta Medical and Graphic film processor (Model SRX-101A). The membrane was subsequently rehydrated in TBS-T and used for Western blot analysis. Specifically, the membrane was probed with anti-human LARP1 rabbit polyclonal antibody (AbCam, catalogue no. ab86359) in TBS-T for 1 h at room temperature on an orbital shaker, washed twice (5 min each time) with TBS-T and incubated with anti-rabbit horseradish peroxidase (HRP)-conjugated IgG (Millipore Sigma, catalogue no. A0545) for 1h at room temperature on an orbital shaker, at which point the unbound secondary antibody was washed three times in TBS-T (5 min each time). Protein was detected by enhanced chemiluminescence (ECL) using the Western Lightning Plus-ECL reagent (Perkin Elmer Inc., catalogue no. NEL105001EA). ECL signal was detected by autoradiography using HyBlot CL film (Denville Scientific Inc., catalogue no. E3018). Samples of lysates (5 μl) were also analyzed by SDS-PAGE/Western blot for mTORC1 activation using phospho-specific antibodies against T389 on S6K1 and T37/T46 on 4E-BP1 as described in the *SDS-PAGE/Western blot* experimental procedures section.

### Structural modeling

The *Arabidopsis thaliana* RAPTOR structure complexed with the *Homo sapiens* PRAS40 TOS peptide shown in **Fig. 2B** was obtained from the PDB, 5WBL (65). Images were prepared using PyMOL 2.2.0 (Schrödinger, LLC, 2015). *Homo sapiens* LARP2 protein sequence (NP_060548) was used to obtain a 3D model of its DM15 domain using Modeller 9.19 (homology modelling) (Webb and Sali, 2016) (**Fig. 2F** and **Fig. 2I**) and the human LARP1 structures obtained from the Protein Data Bank (PDB) accession codes 5v4r (m^7^GTP-bound) and 5v7c (5'TOP RNA-bound) (Lahr *et al*., 2017) were used as template. The DM15 domains of LARP1 and LARP2 are 87% identical and 93% similar based on amino acid sequence alignment (comprising residues 695 to 845 of LARP2). Modeller DOPE score was used to rank models and the best predicted model was selected for further refinement. Coot (version 0.8.6.1, (Emsley *et al*., 2010)) was used to manually refine the model, fix steric clashes, improper angles and other modeling issues. Prosa-web (Wiederstein and Sippl, 2007) and Ramachandran plot analysis module of Coot was used to validate the quality of the model, with 98.7% of the residues in favored regions and no outliers.

### Mass spectrometry

The raw data for the rapamycin screen were extracted from the previous study (47). The raw MS/MS spectra were searched against a composite database of the mouse IPI protein database and its reversed complement using the Sequest algorithm. Search parameters include a static modification of 57.02146 Da for Cys, and a dynamic modification of phosphorylation (79.96633 Da) on Ser, Thr and Tyr. Furthermore, a dynamic modification was also considered for oxidation (15.99491 Da) on Met, and stable isotope (10.00827 Da) and (8.01420 Da) on Arg and Lys, respectively. Search results were filtered to a 1% false-discovery rate (FDR) using the linear discriminator function (78). Phosphorylation site localization was assessed by the ModScore algorithm and peptide quantification was performed by using the CoreQuant algorithm (78).

### RNA-Electrophoretic Mobility Shift Assay (RNA-EMSA)

RNA-EMSAs were performed and imaged as reported previously using the same amount of RNA (≤200 pM) regardless of labeling efficiency (21). 5x protein stocks were prepared in protein dilution buffer (50 mM Tris-HCl, pH 7.5, 250 mM NaCl, 25% glycerol, 2 mM DTT). 10 μL reactions contained 2 μL 5x protein stock, 2 μL 5x RNA stock at 20 nM, 2 μL 5x binding buffer, resulting in final concentrations of 20 mM Tris-HCl, pH 8, 150 mM NaCl, 10% glycerol, 1 mM DTT, 0.5 μg tRNA and 1 μg BSA. Reactions were incubated on ice for 30 minutes and 8 μL were loaded on 7–8% polyacrylamide (29:1) native 0.5x TBE gels at 4°C. Gels were run at 120 V for 40 minutes, dried, and exposed overnight. Phosphor screens (GE Lifesciences) were imaged on a Typhoon FLA plate reader (GE Lifesciences) and quantitated using Imagequant TL (GE Lifesciences). All RNAs were snap-cooled by heating at 95°C in 1X binding buffer for 1 minute and immediately transferred to ice for 20 min.

### Protein expression and purification

Plasmids expressing mutants of DM15 were generated using site-directed mutagenesis and confirmed by sequencing. The LARP1 coding sequence (amino acids 796–946 from LARP1) was cloned by PCR from full-length LARP1 coding sequence into a modified pET28a vector. The resulting construct expressed DM15 with an N-terminal His6-MBP tag followed by a tobacco etch protease cleavage site and glycine6 linker. This expression plasmid was transformed into BL21(DE3) *E. coli* cells and grown overnight on LB agar plates supplemented with 30 μg/ml kanamycin. The His6-MBP-DM15 fusion protein was expressed by autoinduction for 3 h at 37°C and at 18°C for 18 h. Cells were collected by centrifugation, flash frozen in liquid nitrogen, and stored at −80°C until used. 2g of cells were resuspended by gentle stirring in NiNTA lysis buffer (50 mM Tris-HCl, pH 7.5, 400 mM NaCl, 10 mM imidazole, 10% glycerol) for 1 h with protease inhibitor cocktail (PMSF, Leupeptin, Bestatin and Aprotinin). Cells were lysed by homogenization and clarified by centrifugation at 12,000 RPM at 4°C for 30 minutes. The soluble fraction was nutated with 4 mL HisPur Ni-NTA Resin (ThermoFisher product 88221) for 2 h at 4°C. The beads were washed 2 times in 50 mL lysis buffer and 3 times with wash buffer (50 mM Tris-HCl, pH 7.5, 400 mM NaCl, 35 mM imidazole, 10% glycerol). His6-MBP-(665–947) fusion protein was eluted from beads in 15 mL elution buffer (50 mM Tris pH 7.5, 400 mM NaCl, 250 mM imidazole, 10% glycerol). The N-terminal His6-MBP tag was removed by the addition of 2 mg tobacco etch protease for cleavage overnight. Cleaved DM15 protein was further purified of RNA and protein contaminants by tandem HiTrap Q and HiTrap SP columns (GE Lifesciences). DM15 protein free of nucleic acid contaminants was eluted from the HiTrap SP column with gradient from 150 mM NaCl to 1 M NaCl over 50 mL. MBP flowed through both columns while untagged DM15 eluted at 35% elution buffer. Any uncleaved fusion protein eluted at 20% elution buffer, allowing for efficient separation of cleaved DM15. Fractions containing DM15 were pooled and brought to 1 M ammonium sulfate by the dropwise addition of 3M ammonium sulfate with gentle swirling. The protein was diluted to 40 mL in 50 mM Tris-HCl, pH 7, 1 M ammonium sulfate and loaded onto a 5 mL Butyl HP column (GE Lifesciences) at 0.5 mL/min. The Butyl HP column was eluted over 5 CV in 50 mM Tris-HCl, pH 7, 2 mM DTT. The fractions containing DM15 were collected, concentrated, and buffer exchanged with a 10K MWCO Amicon Ultra spin concentrator (Millipore) in 25 mM Tris-HCl, pH 7.5, 2 mM DTT, 250 mM NaCl, and loaded on an equilibrated GE HiLoad 16/600 Superdex 75 gel filtration column run at 1mL/min. DM15-containing fractions were pooled and glycerol increased to 20% before being flash frozen in 10 μL aliquots and stored at −80°C for further use.

### Statistical analysis

All data shown were derived from two biological replicates. Error bars shown in figures 5E-F denote standard deviations (SD) for three independent technical replicates. Error bars shown in figure 5G denote propagated standard deviation (PSD) for each of the total of six replicates in figures 5E-F. Statistical analyses were performed using Prism 5 (GraphPad Software, California). Where indicated, one-way analysis of variance (ANOVA) with Dunnett’s post-hoc tests were performed. Legend: *p<0.05, **p<0.01, ***p<0.001, ****p<0.0001, NS denotes non-significant.

## Supporting information

## Acknowledgments

B.D.F. wishes to dedicate this manuscript to the loving memory of Maria José Morais Fernandes. B.D.F. gratefully acknowledges the financial support received from a Movember Discovery Grant (D2015-02) awarded by Prostate Cancer Canada (PCC) to T.A. and B.D.F. and from a Research Scholar Grant (RSG-17-197-01-RMC) awarded by the American Cancer Society (ACS) to A.J.B. and B.D.F. Y.Y. acknowledges the financial support received from the National Institutes of Health (NIH) (GM114160 and GM122932) and from American Cancer Society (RSG-15-062-01-TBE). C.K.D. acknowledges the financial support received from the Danish Council for Independent Research (6108-00197B). A.J.B. acknowledges the financial support received from the NIH (GM116889) and the ACS (RSG-17-197-01-RMC). T.A. acknowledges the financial support received from PCC Movember Discovery Grant (D2015-02), the Terry Fox Research Institute (TFRI), and the Natural Science and Engineering Research Council (NSERC) for this project.

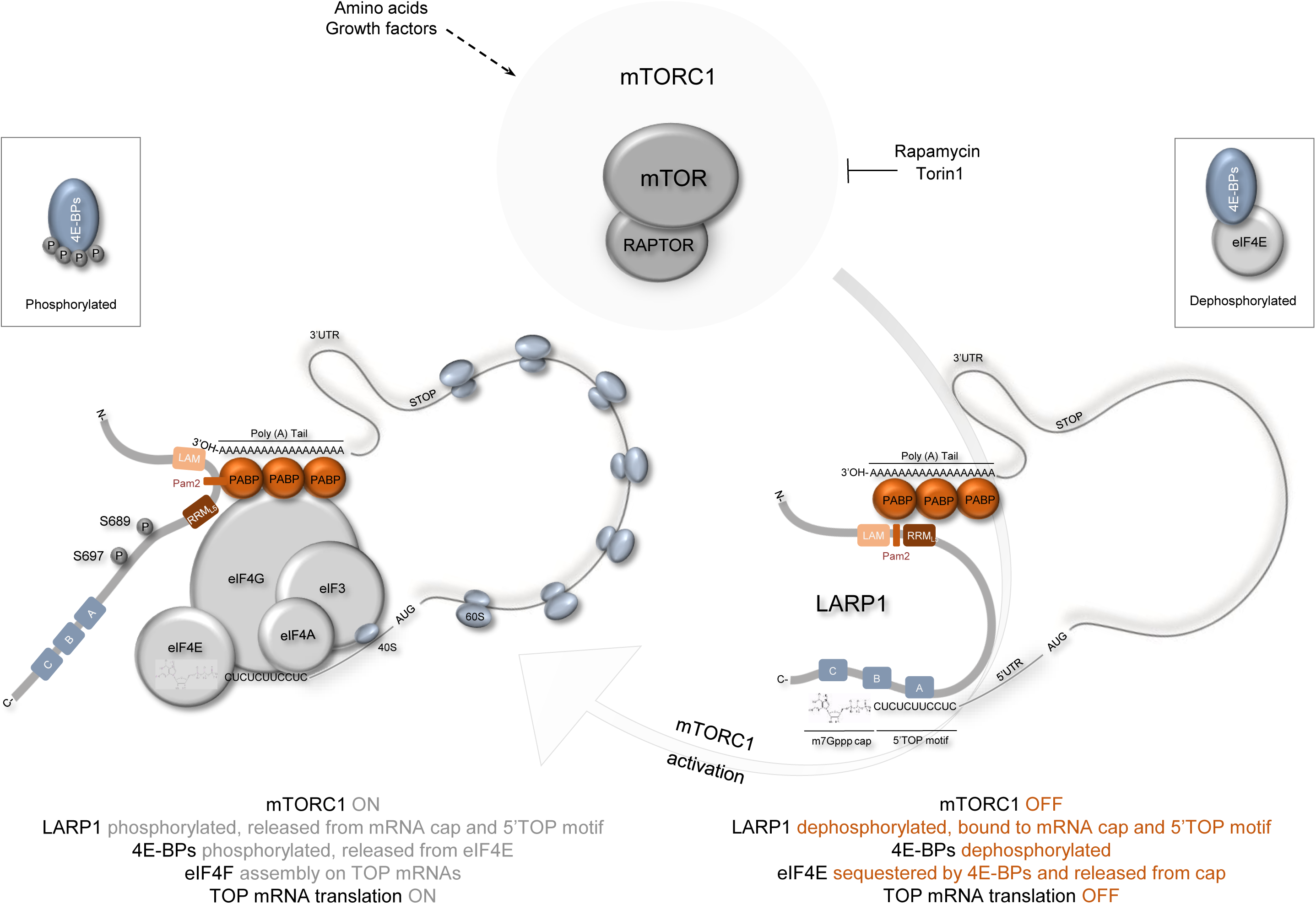
Graphical Abstract. Diagram depicting putative model for mTORC1-mediated release of LARP1 for the N7-methyl guanosine triphosphate (m7Gppp) cap structure and adjacent 5’ terminal oligopyrimidine (5’TOP) motif. mTORC1 phosphorylates S689 and S697 in a rapamycin-dependent manner. Phosphorylation of S689, in particular, contributes to the release of LARP1 from the 5’UTR of RPS6 mRNA, thus allowing for the assembly of the eIF4F complex on 5’TOP mRNAs. Concurrently, mTORC1 phosphorylates multiple serine and threonine residues on 4E-BPs, releasing the latter protein from eIF4E and allowing for the association of eIF4E with eIF4G which together with eIF4A and eIF3 recruit the 40S subunit of the ribosome to TOP mRNAs for translation initiation.

## Footnotes

1 The biochemical significance of each of these mTORC1-mediated phosphorylation events is reviewed in detail in reference number 25. Fonseca, B.D., Graber, T.E., Hoang, H.D., González, A., Soukas, A.A., Hernández, G., Alain, T., Swift, S.L., Weisman, R., Meyer, C., Robaglia, C., Avruch, J. and Hall, M.N. (2016), *Evolution of the Protein Synthesis Machinery and Its Regulation*. Springer International Publishing, pp. 327–411.

2 Percentage identity and similarity between human LARP1 (NP_ 056130) and LARP2 (NP_060548) was determined using the LALIGN protein alignment tool using the following default pairwise alignment options. MATRIX (BLOSUM50), GAP OPEN (−12), GAP EXTEND (−2), E (number) THRESHOLD (10.0), OUTPUT FORMAT (MARKX 0). LALIGN software tool can be found at: https://www.ebi.ac.uk/Tools/psa/lalign/

3 Details of 5’UTR for each of these mRNAs can be found in **Supplemental Fig. 4A**

4 As observed for endogenous LARP1, exogenous (FLAG-tagged) LARP1 also interacts with TOP mRNAs (**Supplemental Fig. 4B and 4C**). In addition, exogenous FLAG-tagged LARP1 also interacts with non-TOP mRNAs; this is especially obvious when FLAG-LARP1 IP data is normalized to input (**Supplemental Fig. 4C-4E**). We speculate that this may be likely the case because endogenous LARP1 is already bound to TOP mRNAs; so proportionally FLAG-LARP1 will bind more efficiently to non-TOP mRNAs (than endogenous LARP1) because these mRNAs are simply freely available for binding. Care must be therefore be taken when interpreting mRNA-binding data obtained from overexpression experiments.

5 RNAse A was reconstituted in 10 mM Tris-HCl (pH 7.5), 15 mM NaCl and 50 % (v/v) glycerol to a final concentration of 10 mg/ml and heated at 96°C for 15 min to inactivate contaminating DNases and cooled down slowly to room temperature. A range of RNAse A concentrations (1, 10 or 100 μg/mL) can be used in the extraction buffer to digest RNA and enhance the interaction between endogenous LARP1 and RAPTOR proteins. A final concentration of 10 μg/mL RNase A in the extraction buffer is recommended for optimal interaction of RAPTOR with LARP1 (**Fig. 3C**). RNAse A was omitted from the extraction buffer for RNA immunoprecipitation experiments.

