## Supplementary material for "LARP1 is a major phosphorylation substrate of mTORC1"

**CLUSTAL O(1.2.4) Human LARP family amino acid sequence alignment**

```
Human LARP1      MLWRVLLSKRPPFPHPELDFQEAPIPSCPGRLPGRKNSVALAAAPRKEPTGDREKPLPFP
Human LARP2      ------------------------------------------------------------
Human LARP2ΔCT   ------------------------------------------------------------
Human La         ------------------------------------------------------------
Human LARP4      ------------------------------------------------------------
Human LARP5      ------------------------------------------------------------
Human LARP6      ------------------------------------------------------------
Human LARP7      ------------------------------------------------------------


Human LARP1      VLAPFSNPEHSAPAKVVRAAVPKQRKGSKVGDFGDAINWPTPGEIAHKSVQPQSHKPQPT
Human LARP2      -----------------------------------MENWPTPSELVNTGFQSVLSQGNKK
Human LARP2ΔCT   -----------------------------------MENWPTPSELVNTGFQSVLSQGNKK
Human La         ------------------------------------------------------------
Human LARP4      ------------------------------------------------------------
Human LARP5      ------------------------------------------------------------
Human LARP6      ------------------------------------------------------------
Human LARP7      ------------------------------------------------------------


Human LARP1      RKLPPKKDMKEQEKGEGSDSKESPKTKSDESGEEKNGDEDCQRGGQKKKGNKHKWVPLQI
Human LARP2      PQN-RKEKEEKVEKRSNSDSKENRETKLNGPGEN-VSEDEAQSSNQRKRANKHKWVPLHL
Human LARP2ΔCT   PQN-RKEKEEKVEKRSNSDSKENRETKLNGPGEN-VSEDEAQSSNQRKRANKHKWVPLHL
Human La         ------------------------------------------------------------
Human LARP4      ------------------------------------------------------------
Human LARP5      -------------MTSDQDAKV-------------VAEPQTQRVQEGKDSAHLMNGPISQ
Human LARP6      ---------------------------------------MAQSGGE--------------
Human LARP7      ------------------------------------------------------------


Human LARP1      DMKPEVPREKLASRPTRPPEPRHIPANRGEIKGSESATYVPVAPPTPAWQPEI-------
Human LARP2      DVVRSESQERPGSRNSSRCQPEANKP-----------THNNRRNDTRSWKRDR-------
Human LARP2ΔCT   DVVRSESQERPGSRNSSRCQPEANKP-----------THNNRRNDTRSWKRDR-------
Human La         ------------------------------------------------------------
Human LARP4      -----------------MLLFVEQVA----------SKGTGLNPNAKVWQEIAPGNTDAT
Human LARP5      ----------TTSQTSSIPPLSQVPA----------TKVSELNPNAEVWGAPVLH-LEAS
Human LARP6      ----------------ARPGPKT------------------------AVQ----------
Human LARP7      ------------------------------------------------------------


Human LARP1      ----KPEPAWHDQDETSSVKSDGAGGARASFRGRGRGRGRGRGRGRGGTRTHFDYQFGYR
Human LARP2      ----E---KRDDQDDVSSVRSE-GGNIRGSFRGRGRGRGRGRGRGRGNPRLNFDYSYGYQ
Human LARP2ΔCT   ----E---KRDDQDDVSSVRSE-GGNIRGSFRGRGRGRGRGRGRGRGNPRLNFDYSYGYQ
Human La         ------------------------------------------------------------
Human LARP4      PVTHGTESSWHEIAATSGAHP-E-G---------------NAELSE-DICKE--YEVMYS
Human LARP5      SAADGVSAAWEEVAGHHADRGPQ-G---------------SDANGDGDQGHE--NAALP-
Human LARP6      -----IRVAIQEAEDVDELEDEEEG---------------AETRGAGDPARYLSPGWGSA
Human LARP7      ------------------------------------------------------------

Human LARP1      KFD--------------GV-EGPRTPKYMNNITYYFDNVSSTELYSVDQELLKDYIKRQI
Human LARP2      EHG--------------ERTDQPFQTELNTSMMYYYDDGTGVQVYPVEEALLKEYIKRQI
Human LARP2ΔCT   EHG--------------ERTDQPFQTELNTSMMYYYDDGTGVQVYPVEEALLKEYIKRQI
Human La         --------------------------------------MAE-NGDNEKMAALEAKICHQI
Human LARP4      SSCETTRNTTGIEESTDGMILGPEDLS---YQ--IYD-VSGESNSAVSTEDLKECLKKQL
Human LARP5      D------PQESDPADMNALALGPSEYD---SLPENSE-TGGNESQPDSQEDPREVLKKTL
Human LARP6      SEE--------------EPSRGHSG-TT-ASGGENEREDLE-QEWKPPDEELIKKLVDQI
Human LARP7      --M--------------ETESGNQE-KVMEEESTEKKKEVE-KKKRSRVKQVLADIAKQV
                                                         .            :   :

Human LARP1      EYYFSVDNLERDFFLRRKMD--ADGFLPITLIASFHRVQALTTDISLIFAALKDSKVVEI
Human LARP2      EYYFSVENLERDFFLRGKMD--EQGFLPISLIAGFQRVQALTTNLNLILEALKDSTEVEI
Human LARP2ΔCT   EYYFSVENLERDFFLRGKMD--EQGFLPISLIAGFQRVQALTTNLNLILEALKDSTEVEI
Human La         EYYFGDFNLPRDKFLKEQIK-LDEGWVPLEIMIKFNRLNRLTTDFNVIVEALSKSKAELM
Human LARP4      EFCFSRENLSKDLYLISQMD--SDQFIPIWTVANMEEIKKLTTDPDLILEVLRSSPMVQV
Human LARP5      EFCLSRENLASDMYLISQMD--SDQYVPITTVANLDHIKKLSTDVDLIVEVLRSLPLVQV
Human LARP6      EFYFSDENLEKDAFLLKHVRRNKLGYVSVKLLTSFKKVKHLTRDWRTTAHALKYSVVL—
Human LARP7      DFWFGDANLHKDRFLREQIEKSRDGYVDISLLVSFNKMKKLTTDGKLIARALRSSAVVEL
                 :: :.  **  * :*  ::      :: :  :  :..:: *: :      .*

Human LARP1      --VDEKVRRREE---------PEKWPLPP--IVDYSQTDFSQ------------------
Human LARP2      --VDEKMRKKIE---------PEKWPIPGPPPRSVPPTDFSQ------------------
Human LARP2ΔCT   --VDEKMRKKIE---------PEKWPIPGPPPRSVPPTDFSQ------------------
Human La         EISEDKTKIRRS---------PSK-PLP--EVTDEYKNDVKNRSVYIK------GFP---
Human LARP4      DEKGEKVRPSHKRCIVILREIPETTPIE--EVKGLFKSENCP------------------
Human LARP5      DEKGEKVRPNQNRCIVILREISESTPVE--EVEALFKGDNLP------------------
Human LARP6      ELNEDHRKVRRT---------TPV-PLF--PNENLPSKMLLVYDLYLSPKLWALATPQKN
Human LARP7      DL--EGTRIRRK-----------K-PLG--E----RPKDEDERTVYVE------LLP---
                     :  :                 *:

Human LARP1      --------------LLNCPEFVPRQHYQKETESAPGSPRAVTPVPT-KTEEVSNLKTLPK
Human LARP2      --------------LIDCPEFVPGQAFCSHTESAPNSPRIGSPLSPKKNSETSILQAMSR
Human LARP2ΔCT   --------------LIDCPEFVPGQAFCSHTGNIFSSRANDVTVGYLVQ--FTYLFIYF-
Human La         ---TDATLDDIKEWLEDKGQVLNIQM-----------RRT---L-HKAFKGSIFVVFDSI
Human LARP4a     -KVISCEFAHNSNWYITFQ---------------------------------------SD
Human LARP4b     -KFINCEFAYNDNWFITFE---------------------------------------TE
Human LARP6      GRVQEKVMEHLLKLFGTFGVISSVRILKPGRELPPDIRRISSRYSQVGTQECAIVEFEEV
Human LARP7      ---KNVNHSWIERVFGKCGNVVYISI-----------PHY---KSTGDPKGFAFVEFETK


Human LARP1      GLSAS------LPDLDSE--N-WIEVKKRPRPSPARPKKSEESRFSHLTSLPQQLPSQQL
Human LARP2      GLSTS------LPDLDSE--P-WIEVKKRHQPAPVKLRESVSV--------PE-GSLNQL
Human LARP2ΔCT   --------------------Q-FI------------------------------------
Human La         ESAKK---FVETPGQKYKE-TDLLIL-FKDDYF-A--KKNEERKQNKV------------
Human LARP4      TDAQQAFKYLREEVK-----------TFQGKPIMARIK----A---INTFFAKNGY-RLM
Human LARP5      ADAQQAYKYLREEVK-----------TFQGKPIKARIKAKAIA---INTFLPKNGF-RPL
Human LARP6      EAAIKAHEFMITESQGKENMKAVLIGMKPPKKKPAKDKNHDEEP----T-----AS-IHL
Human LARP7      EQAAKAIEFLNNPPEEAPRKPGIFPKTVKNKPIPALRVVEEKKKKKKKK-----GR-MKK
```

```
Human LARP1       MSKDQDE----QEELDFLFDEEMEQMDGRKN--------TFTAWSDEESDYEI-------
Human LARP2       CSSEEPE----QEELDFLFDEEIEQI-GRKN--------TFTDWSDNDSDYEI-------
Human LARP2ΔCT    ------------------------------------------------------------
Human La          EAKLRAK----QEQEA-------------KQ--------KLE----EDAEMKS-------
Human LARP4       DSSIYSHPIQTQAQYASPV--FMQPVYNPHQQYSVYSIV-PQSWSPNPTPYFETPLAPFP
Human LARP5       DVSLYA-----QQRYATSF--YFPPMYSPQQQFPLYSLITPQTWSATHSYLDPPLVTPFP
Human LARP6       NKSLNKR----VEELQ--------YM-GD-----------------ESSANSS-------
Human LARP7       EDNIQAK----EENMDTS-NTSISKM-KRSR--------PTS----EGSDIES-------

Human LARP1       ----------------DDRDVNKILIVTQTP-------H----YMRRHPGGDR-------
Human LARP2       ----------------DDQDLNKILIVTQTP-------P----YVKKHPGGDR-------
Human LARP2ΔCT    ------------------------------------------------------------
Human La          ----------------LEEKIGCLLKFSG--------------DLDDQTCREDLHILFSN
Human LARP4       NGSFVNGFNSPGSYKTNAAAMNMGRPFQK---N------RVKPQFR-SSGGSE-HSTEGS
Human LARP5       NTGFINGFTSPAFKPAASP-LTSLRQYPPRSRN------PSKSHLR-HA--------IPS
Human LARP6       ----------------SDPESNPTSPMAGRRHAATNKLS---------PSGHQNLFLSPN
Human LARP7       ----------------TEPQKQCSKKKKKRDRVEASSLPEVRTGKRKRSSSEDAESLAPR

Human LARP1       ---------------------TGNHT--SRAKMSAELAKVINDGLFYY--EQDLWAEK-F
Human LARP2       ---------------------TGTHM--SRAKITSELAKVINDGLYYY--EQDLWMEE-D
Human LARP2ΔCT    ------------------------------------------------------------
Human La          H----GEIKW---IDFVRGAKEGIILFKEKAKEALGKAKDANNGNLQLRNKEVTWEVLEG
Human LARP4       VSLGDGQLNRYSSRNFPAERHNPTVT-GHQ-------------EQTYLQKETS-----TL
Human LARP5       AERGPGLLESPSIFNFTADRLINGVR-SPQTRQAGQTRTRIQNPSAYAKREAGPGRVEPG
Human LARP6       AS--PCTSPW---SSPLAQR-----------KGVSRKSPLAEEGRLNCSTSPEIFRKCMD
Human LARP7       -------------------SKVKKIIQKDIIKEASEASKE--NRDIEISTEE---EKDTG

Human LARP1       EPEYSQIKQEVENFKK--------------VNMISREQFDTLTPEPPVDPNQEVPPGPPR
Human LARP2       ENKHTAIKQEVENFKK--------------LNLISKEQFENLTPELPFEPNQEVPVAPSQ
Human LARP2ΔCT    ------------------------------------------------------------
Human La          EVEKEALK------------------KIIEDQQESLNKWKSKGRRFK--GKG---KGNKA
Human LARP4       QVEQNGDY--GRGRRTLFRGRRRREDDRISRPHPSTAESKAPTPKFDLLA-SNFPPLPGS
Human LARP5       SLESSPGL--GRGRKNSFGYRKKREEKFTSSQTQSPTPPKPPSPSFELGL-SSFPPLPGA
Human LARP6       YSSDSS-------------------------VTPSGSPWVRRRRQA--------------
Human LARP7       DLKDSSLLKTKRKHKKKHKERHKMGEEVIPLRVLSKSEWMDLKKEYL--ALQ---KASMA

Human LARP1       FQQV------------PTDALANKLFGAPE---PSTIARSLPTTVPESPNYRNTRTPR-T
Human LARP2       SRQGGVQGVLHIPKKDLTDELAQKLFDVSEI-TSAAMVHSLPTAVPESPRIHPTRTPK-T
Human LARP2ΔCT    ------------------------------------------------------------
Human La          AQPGSGKGKVQFQGKK------TKFASDDE------HDEHDENGATGPVKRAREETDKEE
Human LARP4       SS--RMPGELVLEN-----RMSDVVKGVYKEKDNEEL--TISCPVPADEQTECTSAQQLN
Human LARP5       AG--NLKTEDLFEN-----RLSSLIIGPSKERTLSAD--ASVNTLPVVVSR--------E
Human LARP6       ------------------------------------------------------------
Human LARP7       SL----------KKTI------SQIKSESE------M—ETDSGVPQNTGMKNEKTANRE

Human LARP1       P----RTP-QLKDSSQTSRFYPVVKEGRTLDAKMPRKRKTRHSSNPPLESHVGWVMDSRE
Human LARP2       P----RTP-RLQDPNKTPRFYPVVKEPKAIDVKSPRKRKTRHSTNPPLECHVGWVMDSRD
Human LARP2ΔCT    ------------------------------------------------------------
Human La          P----ASKQQKTENGAGDQ-----------------------------------------
Human LARP4       MSTSSPCAAELTALSTTQQEKDLIEDSS-----VQKDGLNQTT--------IPVSPPSTT
Human LARP5       PSVPASCAVSATYE-RSPSPAHLPDDPK-----VAEKQRETHS---------------VD
Human LARP6       -----EMGTQEKSPGTSPLLSRKMQTADGL----------------------PVGVLRLP
Human LARP7       -----ECRTQEKVNATGPQFV------SGV----------------------IVKIISTE

Human LARP1       HRPRTASIS---SSPS---EG---TPTVGSYGCTPQSLPKFQHPSHELLKENGFTQHVYH
Human LARP2       RGPGTSSVSTSNASPS---EG---APLAGSYGCTPHSFPKFQHPSHELLKENGFTQQVYH
Human LARP2ΔCT    ------------------------------------------------------------
Human La          ------------------------------------------------------------
Human LARP4       KPSRASTASPCNNNINAATAVALQEPRKLSYAEVCQ--KPPKEPSSVLV----QPLRELR
Human LARP5       RLPSALTATACKSVQ---VNGAATELRKPSYAEICQ--RTSKEPPSSPL----QPQKEQK
Human LARP6       RGPD----------------------------NT----RGFHGHERSRA-----------
Human LARP7       PLPGRKQVRDTLAAIS---EVLYVDLLEGDTECH----ARFKTPEDAQAVINAYTE----

Human LARP1       KYRRRCLNERKRLGIGQSQEMNTLFRFWSFFLRDHFNKKMYEEFKQLALE---DAKEGYR
Human LARP2       KYRRRCLSERKRLGIGQSQEMNTLFRFWSFFLRDHFNKKMYEEFRQLAWE---DAKENYR
Human LARP2ΔCT    ------------------------------------------------------------
Human La          ------------------------------------------------------------
Human LARP4       SNVVSPTKNEDNGAPENSVEK---------------------------------------
Human LARP5       PNTVGCGKEEKKL------AE---------------------------------------
Human LARP6       -----CV-----------------------------------------------------
Human LARP7       INKKHCWKLEIL-------SGDHEQRYWQKILVDRQAKLNQPREKKRGTEKLITKAEKIR

Human LARP1       Y----GLECLFRYYSYGLEKKFRLDIFKDFQEETVKDYEAGQLYGLEKFWAFLKYSKAKN
Human LARP2       Y----GLECLFRFYSYGLEKKFRREIFQDFQEETKKDYESGQLYGLEKFWAYLKYSQSKT
Human LARP2ΔCT    ------------------------------------------------------------
Human La          ------------------------------------------------------------
Human LARP4       ------------------------------------------------------------
Human LARP5       ------------------------------------------------------------
Human LARP6       ------------------------------------------------------------
Human LARP7       LAKTQQASKHIRFSEYD-------------------------------------------

Human LARP1       LDIDPKLQEYLGKFRRLEDFRVDPPMGEEGNHKRHSVVAGGGGGEGRKRCPSQSSSRPAA
Human LARP2       QSIDPKLQEYLCSFKRLEDFRVDPPISDEFGRKRHSSTSGEES--NRHRLPPNSSTKPPN
Human LARP2ΔCT    ------------------------------------------------------------
Human La          ------------------------------------------------------------
Human LARP4       ----PHEKPEARASKDYSGFRGNIIPRGAAGKIREQ------------RRQFSHRAIPQG
Human LARP5       ----PA-----------ERYREPPALKSTPGAPRDQ------------RRPAGGRPSPSA
Human LARP6       ------------------------------------------------------------
Human LARP7       ------------------------------------------------------------

Human LARP1       MISQPPTPPTGQPVREDAKWTSQHSNTQTLGK
Human LARP2       AAKPTSTSELQVPINSPRRNISPESSDNSH—
Human LARP2ΔCT    --------------------------------
Human La          --------------------------------
Human LARP4       VTRR-NGKEQYVPPRSPK--------------
Human LARP5       MGKR-LSREQSTPPKSPQ--------------
Human LARP6       --------------------------------
Human LARP7       --------------------------------
```

**Supplemental Figure 1. Amino acid sequence alignment of human LARP superfamily protein members.**

**Supplemental Figure 2**

A

**Clustal W alignment of LARP1 across eukaryotes**

```
M.musculus        MATQVEPLLPAGAPLLQAEEHGLARKKPAPDAQAESGPGDGGGEPDGGVRRPRPACARPG
H.sapiens         ------------------------------------------MLWRVLLSKRPPFPHPE
G.gallus          ------------------------------------------------------------
X.tropicalis      ----------------------------------MAAQVEAPLLPANPLLPPLQQQPPPE
D.rerio           -----------------------------------------MATEVETLPPNNPLLQPEE


M.musculus        RDGAERESPRPPAAAEAPAGSDGEDGGRRDFVEAPPPKVNPWTKHAPPPAAVNGQPPPEP
H.sapiens         LDFQEAPIPSCPGRLPGRKNSVALAAAPRKEPTGDREKPLPFPVLAP-------FSNPEH
G.gallus          ------------------------------------------------------------
X.tropicalis      EMIEERRPEGGKQRQAKEDGGTGGEEVQLRKEFVEAPPPKVNPWMKAGGSTVNGQLPAEP
D.rerio           PQHKDGEYPLQERHTADSGSPGAGQEEQAAGREISEAPLPKVNPWTKKMSGVNGQPAAEH


M.musculus        SAPAKVVRAAAPKPRKGSKVGDFGDAVNWPTPGEIAHKSVQPQSHKPQPARKLPPKKDMK
H.sapiens         SAPAKVVRAAVPKQRKGSKVGDFGDAINWPTPGEIAHKSVQPQSHKPQPTRKLPPKKDMK
G.gallus          ----------------------------------------------------------MK
X.tropicalis      PVPAKVVKAGNSRPRRGSKVGDFGDATNWPTPGEIAHKTVQPPTKLQPGR-----KSLGK
D.rerio           SGPTKVVKAGNSRLRRGGKVGDFGDTNNWPTPGEIATKEVQVLKKP---------TVKRE
                                                                             :

                                 Cluster I
                                  148 151
M.musculus        EQEKGDGSDSKESPKTKSDESGEEKNGDEDCQRGGQKKKGSKHKWVPLQIDMKPEVPREK
H.sapiens         EQEKGEGSDSKESPKTKSDESGEEKNGDEDCQRGGQKKKGNKHKWVPLQIDMKPEVPREK
G.gallus          EQEKGDGSDGKESLKTKSDESGEEKNGDDDNQKSAQKKKGNKHKWVPLQIDMKSEVPRDK
X.tropicalis      KDMKEESADGKENQRVKSDESSEEKNGDDDGQRSANKKKGNKHKWVPLQIEMKPDGPREK
D.rerio           PKGKRESEESKENLKSKSDDSGEDKNTDEESQRSSHRRRGNKQRWVPLMIEVKAEGPRER
                   . * :. :.**. : ***:*.*:** *:: *:..::::*.*::**** *::*.: **::

                                                                         Cluster
                                                                           247
M.musculus        LASRPTRPQE-PRHTPAVRGEMKGSEPATYMPVSVAPPTPAWQPETKVEPAWHDQDETSS
H.sapiens         LASRPTRPPE-PRHIPANRGEIKGSESATYVPVAPP--TPAWQPEIKPEPAWHDQDETSS
G.gallus          TASRNNRQNEQHRHLPNNRGELKG-----------------WHPDNKHERPWLDHDETSS
X.tropicalis      NASRNNRQNEQQRHPLNRSD------------------PKTMYPDNKSERDWQDFDDTSS
D.rerio           SASRNNTRPYDAHRGTHSSRNGLR----------------DWPSERFEQELKDDHDEVSS
                   *** .      ::                             .:   :    * *:.**

                   II
                  250
M.musculus        VKSDGAGGARASFRGRGRGRGRGRGRGGTRTHFDYQFGYRKFDGTEGPRTHKYMNNIT
H.sapiens         VKSDGAGGARASFRGRGRGRGRGRGRGGTRTHFDYQFGYRKFDGVEGPRTPKYMNNIT
G.gallus          VKSEGG-AVRGAFRGRGRGRGRGRGRG---GHFDYQYGYRKFEGSDGSRTQKYASNIT
X.tropicalis      VKSEGA--IRGGFRGRGRGRGRGRGRGGTRSHFDYPYGYRKFDGSDVTRGPKFLSNIT
D.rerio           VKSDGAPYRAG---ARGRGRGRGRGRGRGHYDYYS-YKGSEGKDGAYAQKFGNSMT
                  **  .     .   .*************    *:**    *:  :* : .   *: ..:*


M.musculus        YYFDNVSSNEIYSMDQELLKDYIKRQIEYYFSVDNLERDFFLRRKMDADGFLPITLIASF
H.sapiens         YYFDNVSSTELYSVDQELLKDYIKRQIEYYFSVDNLERDFFLRRKMDADGFLPITLIASF
G.gallus          YYFDNISSTELYSVDQELLKDYIKRQIEYYFSVDNLERDFFLRRKMDSDGFLPITLIASF
X.tropicalis      YYYDNMSSSELYSVDQELLKDYIKRQIEYYFSVDNLQRDFFLRRKMDSEGFLPVGLIASF
D.rerio           YYYDNMSSAELYSVDQDLLKDYIKRQIEYYFSVDNLERDFFLRRKMDEGGFLPVSLIASF
                  **:**:** *:**:**:*******************:**********  ****: *****

                                                                        PAM2
                                                                        420-
M.musculus        HRVQALTTDISLIFAALKDSKVVEMVEEKVRRREEPEKWPLPGPPIVDYSQTDFSQLLNC
H.sapiens         HRVQALTTDISLIFAALKDSKVVEIVDEKVRRREEPEKWPLP--PIVDYSQTDFSQLLNC
G.gallus          HRVQALTTDISLIIKALKDSKVVEIVDQKIRRKEQPEKWALPGPPMADYTQTDFSQFINC
X.tropicalis      HRVQALTTDIELIVKALKDSKVVEIIDEKIRRKDQPDMWPLPGPSLSDASQTDFNQLINC
D.rerio           HRVQALTTDVSLILQALKDSKVVDIIDMKMRCKEEPQKWPLPDLCLPDQAQTDFAQFIHC
                  *********:.**. ********:::: *:* :::*: *.**   : * :**** *:::*

                   motif                  Cluster III
                   -430     438 440 444 449                    471
M.musculus        PEFVPRQHYQKETESAPGSPRAVTPVP-TKTEEVSNLKTLPKGLSASLPDLDSESWIEVK
H.sapiens         PEFVPRQHYQKETESAPGSPRAVTPVP-TKTEEVSNLKTLPKGLSASLPDLDSENWIEVK
G.gallus          PEFVPRQSFQKETESAPGSPRPTSPVP-TKTEEVSNLKTMPKGLSTSLPDLDSETWIEVK
X.tropicalis      PEFVPRQGFQKETESAPGSPRTSSPARTTKSEEVSNLKTMPKGLSASLPDLDSDNWIEVK
D.rerio           PEFIPR--HLPASPSPAGTPRSGTPRSSTPHQELCSLKMMPKGLSASLPDLDSECWIEVK
                  ***:**  .   : *..*:**. :*   *  :*:..** :*****:*******: *****


M.musculus        KRPRPSPARPKKPEEPRFSHPTALPQQLPSQQLMSKDQDEQEELDFLFDEEMEQMDGRKN
H.sapiens         KRPRPSPARPKKSEESRFSHLTSLPQQLPSQQLMSKDQDEQEELDFLFDEEMEQMDGRKN
G.gallus          KRPRPSPARPKKPEES-----------KTPQQQKQKDQDEPEELDFLFDEEMEQMDGRKN
X.tropicalis      KRPRPSPARTKDEVKS----PQVVTPLQVTPQKPPKDQDEPEELDFMFDEEMEQMDGRKN
D.rerio           KRPRPSPARPKKEDVR---------CVLQMGQVSAEEQEEPEELDFMFDEEMQQMDGRRN
                  *********.*.                   *   ::*:* *****:*****:*****:*

                     Cluster IV
                     550 554 556
M.musculus        TFTAWSEEDSDYEIDDRDVNKILIVTQTPPYMRRHPGGDRTGNHTSRAKMSAELAKVIND
H.sapiens         TFTAWSDEESDYEIDDRDVNKILIVTQTPHYMRRHPGGDRTGNHTSRAKMSAELAKVIND
G.gallus          TFTDWSDEDSDYEIDDRDVNKILIVTQTPPYLRRHPGGDRTGNHTSRAKMSSELAKVIND
X.tropicalis      TFTDWSDEDSDYEIDDRDVNKILIVTQTPPSLRKHPGGDRTGNHTSRAKMSSEFAKVIND
D.rerio           TFTDWSDEDSDCEIDDRDVNKILIVTQTPPYLRKHPGGDRTGHHTSRSKLSSELVKIIND
                  *** **:*:** ****************   :*:********:****:*:*:*:.*:***


M.musculus        GLFYYEQDLWTEKFEPEYSQIKQEVENFKKVNMISREQFDTLTPEPPVDPNQEVPPGPPR
H.sapiens         GLFYYEQDLWAEKFEPEYSQIKQEVENFKKVNMISREQFDTLTPEPPVDPNQEVPPGPPR
G.gallus          GLYYYEQDLWTEQFEPEYSQIKQEVENFKKVNMISREQFDTLTPEPPVDPNQEVPPGPPR
X.tropicalis      GLFYYEQDLWTEQFEPEYSQIKQEVENFKKVNMISREQFDTLTPEPPVDPNQEVPPGPPR
D.rerio           GLFYYEQDLWDDSGEPEYAIIKQEVENFKKVHLISREQFDCLTPEPPVDPNQEVPPGPPR
                  **:******* :. ****: ***********::******* *******************
```

A

```
                                                         Cluster V
                                                   689 692 697 700
M.musculus     FQQVPTDALANKLFGAPEPSTIARSLPTTVPESPNYRNARTPRTPRTPQLKDSSQTPRFY
H.sapiens      FQQVPTDALANKLFGAPEPSTIARSLPTTVPESPNYRNTRTPRTPRTPQLKDSSQTSRFY
G.gallus       FQQVPTDALANKLFGVPEPSTIARSLPTTVPESPNYRNARTPRTPRTPQLKDPTQTPRFY
X.tropicalis   FQQVPTDDLANKLFGAPEPSTIARSLPTTVPESPNYRNARTPRTPRTPRLKDPTLTPRFY
D.rerio        PTQIPTEALANKLFGAPDPSCMARSLPTTVPDSPSHRSARTPRTPRTPRLKD-AQTPRFY
                 *:**: *******.*:** :*********:**.:*.:*********:*** : *.***

                                                          Cluster VI
                                                              768    774
                                                              770    776
                                                  747         772    779
M.musculus     PVVKEGRTLDAKMPRKRKTRHSSNPPLESHVGWVMDSREHRPRTASIS--SSPSEGTPAV
H.sapiens      PVVKEGRTLDAKMPRKRKTRHSSNPPLESHVGWVMDSREHRPRTASIS--SSPSEGTPTV
G.gallus       PVVKEGRTIDAKTPRKRKTRHSSNPPMECHVGWVMDSREHRPRTASIS--SSPSEGTPAV
X.tropicalis   PVVKEGRTIDAKTPRKRKTRHSSNPPMECHVGWVMDSREHRPRTASVSSTASPSEVAPAL
D.rerio        PVVKDGRPLDAKTPRKRKTRHSSNPPMECHVGWVMDSREHRSRTASVSSNASPSEGTAAM
               ****:**.:*** ************:*.************.****:*  :**** :.::

               Cluster VI
                784  788
                785  791
M.musculus     GSYGCTPQSLPKFQHPSHELLKENGFTQHVYHKYRRRCLNERKRLGIGQSQEMNTLFRFW
H.sapiens      GSYGCTPQSLPKFQHPSHELLKENGFTQHVYHKYRRRCLNERKRLGIGQSQEMNTLFRFW
G.gallus       GSYGCTPQSLPKFQHPSHELLKENGFTQHVYHKYRRRCLNERKRLGIGQSQEMNTLFRFW
X.tropicalis   SNYGCTPQSLPKFQHPSHELLKENGFTQHVYHKYRRRCLNERKRLGIGQSQEMNTLFRFW
D.rerio        GNFGCTPQSLPKFQHPSHELLKENGFTQHVYHKYRRRCLNERKRLGIGQSQEMNTLFRFW
               ..:*********************************************************

M.musculus     SFFLRDHFNKKMYEEFKQLALEDAKEGYRYGLECLFRYYSYGLEKKFRLDIFKDFQEETV
H.sapiens      SFFLRDHFNKKMYEEFKQLALEDAKEGYRYGLECLFRYYSYGLEKKFRLDIFKDFQEETV
G.gallus       SFFLRDHFNKKMYEEFKQLAVEDGKEGYRYGLECLFRYYSYGLEKKFRPDIFKDFQEETI
X.tropicalis   SFFLRDHFNKKMYEEFRQLAVEDSKEGYRYGLECLFRYYSYGLEKKFRVDIFRDFQEETR
D.rerio        SFFLRDHFNRKMYEEFRQLSVEDGKEGYRYGLECLFRFYSYGLERKFRPDIFKDFQEETV
               *********:******:**::**.*************:******:*** ***:******

M.musculus     KDYEAGQLYGLEKFWAFLKYSKAKNLDIDPKLQEYLGKFRRLEDFRVDPPMGEEGNHKRH
H.sapiens      KDYEAGQLYGLEKFWAFLKYSKAKNLDIDPKLQEYLGKFRRLEDFRVDPPMGEEGNHKRH
G.gallus       KDYEAGQLYGLEKFWAFLKYSKAKNLDINSKLQEYLSKFKRLEDFRVDPPMGEEGGHKRY
X.tropicalis   RDYETGQLYGLEKFWAFLKYSKAKNLDIDPILQEYLSKFRRLEDFRVDPPMGEDGGSKRR
D.rerio        KDYEAGQLYGLEKFWAFLKYSKTKNMEIEPKLQEHLSKFRRLEDFRVDPPMEDGVRKRHS
               :***:****************:**::*:.  ***:*.**:***********  :    ::

                                  Cluster VII
                              979             994
M.musculus     PVVAGGSG-EGRKRCPSQSSSRPATGISQPPTTPTG-QATREDAKWTSQHSDTLTLRK--
H.sapiens      SVVAGGGGGEGRKRCPSQSSSRPAAMISQPPTPPTG-QPVREDAKWTSQHSNTQTLGK--
G.gallus       PTTSGGDG---RKRYPSQSSSKTVSQCPSSQAHASSSQPAQEDAKNLSQQVPAVESQTVG
X.tropicalis   HSLSTGEG---RRRYPSQSSSRGSGHQTSTQNSAREEAKAQSHPSTATPDSQTPGAGK--
D.rerio        SSGDG------RRRHPSHPGSRSQTQSQLQPSGAAAGATRDDSRAQSQRDARPPK-----
                          *:* **:..*:             .        .        .  .

M.musculus     -
H.sapiens      -
G.gallus       K
X.tropicalis   -
D.rerio        -
```

B

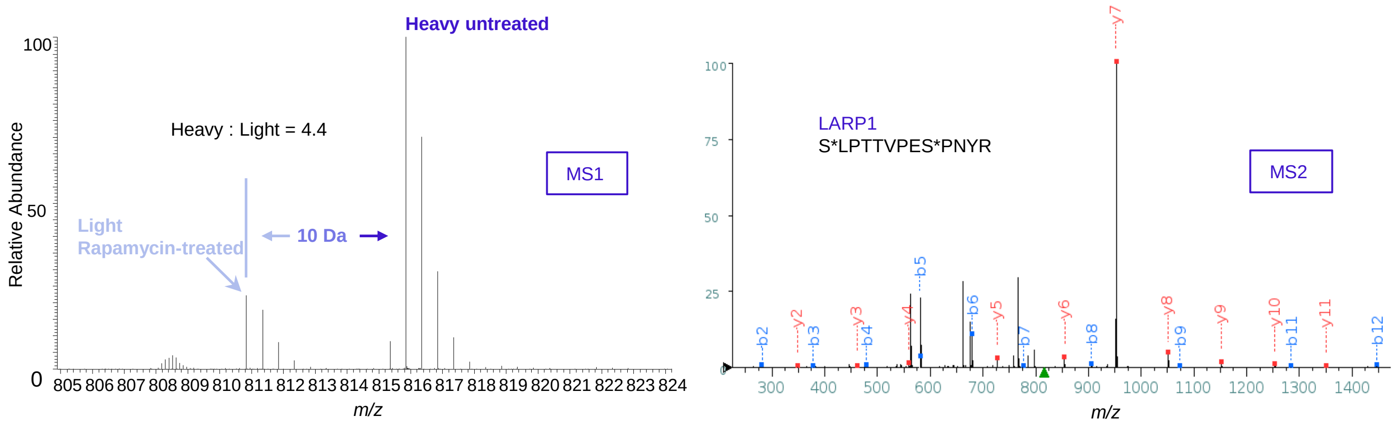


**Supplemental Figure 2. Conservation of rapamycin-sensitive phosphorylation sites on LARP1 and peptide mass/charge fingerprint for S689 and S697 phospho-residues on LARP1.**

**CLUSTAL O(1.2.4) multiple sequence alignment of human LARP1 and human LARP2**

```
Human LARP1      MLWRVLLSKRPPFPHPELDFQEAPIPSCPGRLPGRKNSVALAAAPRKEPTGDREKPLPFP      60
Human LARP2      ------------------------------------------------------------      0


Human LARP1      VLAPFSNPEHSAPAKVVRAAVPKQRKGSKVGDFGDAINWPTPGEIAHKSVQPQSHKPQPT      120
Human LARP2      -----------------------------------MENWPTPSELVNTGFQSVLSQGNKK      25
                                                    *****.*:.:...*    : : .

Human LARP1      RKLPPKKDMKEQEKGEGSDSKESPKTK SDESGEEKNGDEDCQRGGQKKKGNKHKWVPLQI      180
Human LARP2      PQN-RKEKEEKVEKRSNSDSKENRETKLNGPGEN -VSEDEAQSSNQRKRANKHKWVPLHL      83
                 :   *:. :: ** ..*****. :** :  **:  .:::.* ..*:*:.********::

Human LARP1      DMKPEVPREKLASRPTRPPEPRHIPANRGEIKGSESATYVPVAPPTPAWQPEIKPEPAWH      240
Human LARP2      DVVRSESQERPGSRN SSRCQPEANKP-----------THNNRRNDTRSWKRDRE---KRD      129
                 *:  .  :*: .** :   :*.               *:      * :*: : :     .

Human LARP1      DQDETS SVKSDGAGGARASFRGRGRGRGRGRGRGGTRTHFDYQFGYRKFDGV -EGPRT      299
Human LARP2      DQDDVS SVRSE-GGNIRGSFRGRGRGRG RGRGRGRGNPRLNFDYSYGYQEHGERTDQPFQ      188
                 ***:.***:*: .*. *.*****************. * :***.:**::..   : *

Human LARP1      PKYMNNITYYFDNVSSTELYSVDQELLKDYIKRQIEYYFSVDNLERDFFLRRKMDADGFL      359
Human LARP2      TELNTSMMYYYDDGTGVQVYPVEEALLKEYIKRQIEYYFSV ENLERDFFLRGKMDEQGFL      248
                 :  ..: **:*: :..::* *:: ***:************:********* *** :***

Human LARP1      PITLIASFHRVQALTTDISLIFAALKDSKVVEIVDEKVRRREEPEKWPLPP --IVDYSQT      417
Human LARP2      PISLIAGFQRVQALTTNLNLILEALKDSTEVEIVDEKMRKKIEPEKWPIPGPPPR SVPPT      308
                 **:***.*:*******::.**: *****. *******:*:: ******:*     .   *

Human LARP1      DFSQLLNCPEFVPRQHYQKE TESAPGSPRAVTPVPT-KTEEVSNLKTLPKGLSA SLPDLD      476
Human LARP2      DFSQLIDCPEFVPGQAFCSH TESAPNSPRIGSPLSPKKNSETSILQAMSRGLST SLPDLD      368
                 *****::****** * : ..*****.***  :*:   *..*.* *::: :***:******

Human LARP1      SENWIEVKKRPRPSPARPKKSEESRFSHLTSLPQQLPSQQLMSKDQDEQEELDFLFDEEM      536
Human LARP2      SEPWIEVKKRHQPAPVKLRESVSV --------PE-GSLNQLCSSEEPEQEELDFLFDEEI      419
                 ** ******* :*:*.: ::* .          *:    :** *.:: ************:

Human LARP1      EQMDGRKNTFTAW SDEESDYEIDDRDVNKILIVTQTPHYMRRHPGGDRTGNHTSRAKMSA      596
Human LARP2      EQI -GRKNTFTDWSDNDSDYEIDDQDLNKILIVTQTPPYVKKHPGGDRTGTHMSRAKITS      478
                 **: ******* ***::*******:*:********** *:::********.* ****:::

Human LARP1      ELAKVINDGLFYYEQDLWAEKFEPEYSQIKQEVENFKKVNMISREQFDTLTPEPPVDPNQ      656
Human LARP2      ELAKVINDGLYYYEQDLWMEEDENKHTAIKQEV ENFKKLNLISKEQFENLTPELPFEPNQ      538
                 **********:******* *: * ::: **********:*:**:***:.**** *.:***

Human LARP1      EVPPGPPRFQQV ------------PTDALANKLFGAPE --PSTIARSLPTTVPESPNYRN      702
Human LARP2      EVPVAPSQSRQGGVQGVLHIPKKDLTDELAQKLFDVSEITSAAMVH SLPTAVPESPRIHP      598
                 *** .* : :*                 ** **:***.. *   ::::.:****:*****. :

Human LARP1      TRTPRTPRTPQLKDSSQTSRFYPVVKEGRTLDAKMPRKRKTRHS SNPPLESHVGWVMDSR      762
Human LARP2      TRTPKTPRTPRLQDPNKTPRFYPVVKEPKAIDVKSPRKRKTRHS TNPPLECHVGWVMDSR      658
                 ****:*****:*:* .:* ******** :::*.* *********:*****.*********

Human LARP1      EHRPRTASIS---SSPSEGTPTVGSYGCTPQSLPKFQHPSHELLKENGFTQHVYHKYRRR      819
Human LARP2      DRGPGTSSVSTSNASPSEGAPLAGSYGCTPHSFPKFQHPSHELLKENGFTQQVYHKYRRR      718
                 :: * *:*:*   :*****:* .*******:*:*****************:********

Human LARP1      CLNERKRLGIGQSQEMNTLFRFWSFFLRDHFNKKMYEEFKQLALEDAKEGYRYGLECLFR      879
Human LARP2      CLSERKRLGIGQSQEMNTLFRFWSFF LRDHFNKKMYEEFRQLAWEDAKENYRYGLECLFR      778
                 **.********************************:*** *****.**********

Human LARP1      YYSYGLEKKFRLDIFKDFQEETVKDYEAGQLYGLEKFWAFLKYSKAKNLDIDPKLQEYLG      939
Human LARP2      FYSYGLEKKFRREIFQDFQEETKKDYESGQLYGLEKFWAYLKYSQSKTQSIDPKLQEYLC      838
                 :********** :**:****** ****:***********:****::*. .*********

Human LARP1      KFRRLEDFRVDPPMGEEGNHKRHSVVAGGGGGEGRKRCP SQSSSRPAAMISQPPTPPTGQ      999
Human LARP2      SFKRLEDFRVDPPISDEFGRKRHSSTSGEES --NRHRLPPNSSTKPPNAAKPTS TSELQV      896
                 .*:**********:.:* .:**** .:*  .  .*:* * :**::*    .   *

Human LARP1      PVREDAKWTSQHSNTQTLGK     1019
Human LARP2      PINSPRRNISPESSDNSH --    914
                 *:..  :  * .*. ::
```

**Supplemental Figure 3. Alignment of human LARP1 and human LARP2 proteins.**

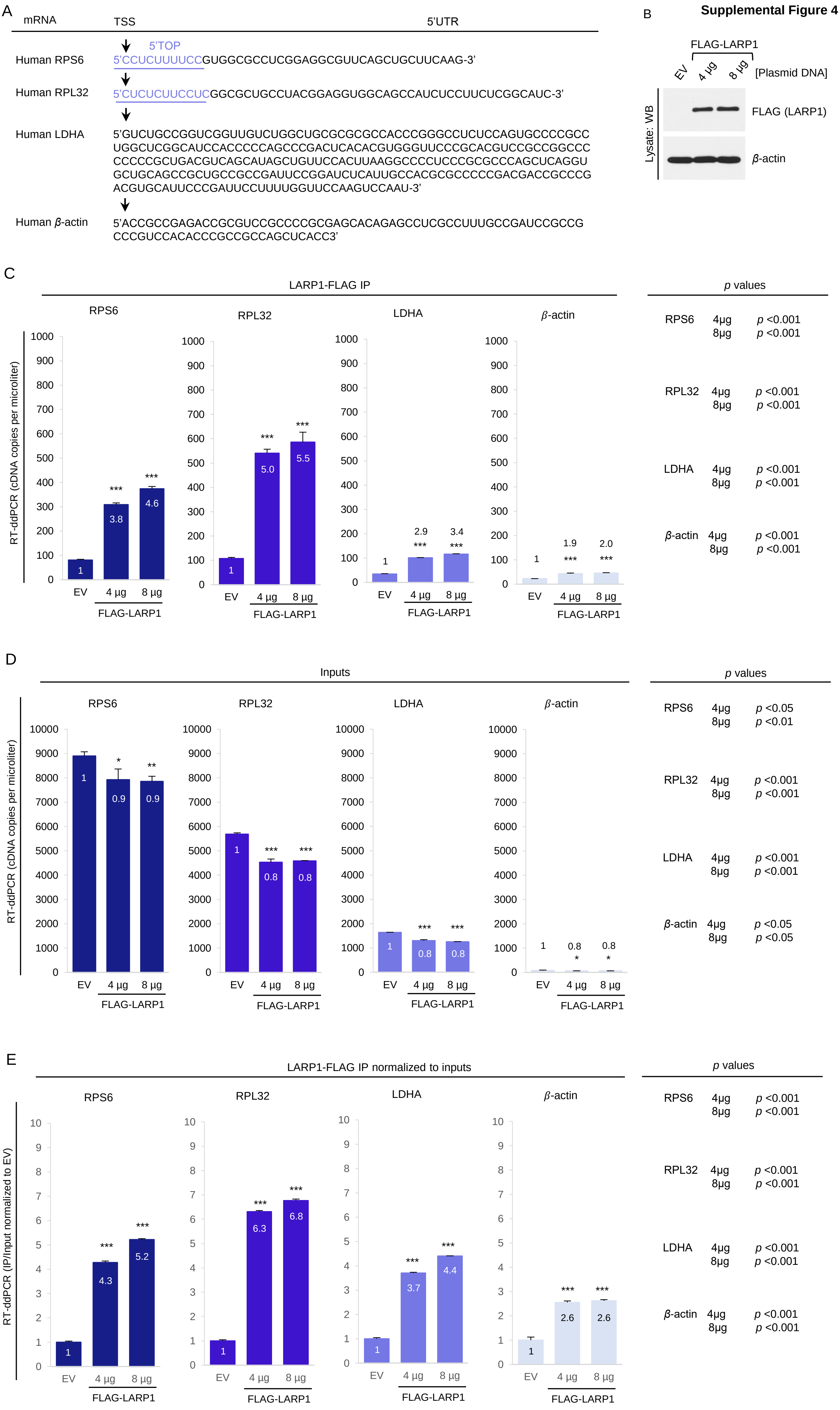


**Supplemental Figure 4. Ectopic LARP1 binds TOP and non-TOP mRNAs.** (A) Complete 5'UTR of ribosomal protein S6 (RPS6), of ribosomal protein L32 (RPL32) and β-actin. Arrow marks the transcriptional start site (TSS) and blue sequence in RPS6 and RPL32 denotes the 5'terminal oligopyrimidine (5'TOP) motif on these transcripts. (B) HEK293T cells were transiently transfected with the indicated amouns of FLAG-tagged LARP1 or control empty vector (EV). Cells were lysed and lysates analyzed for protein expression level by western blot. (C-E) Samples of lysates were used for used for RNA-immunoprecipitation with anti-FLAG antibody. Absolute mRNA abundance was detected by ddPCR.

A

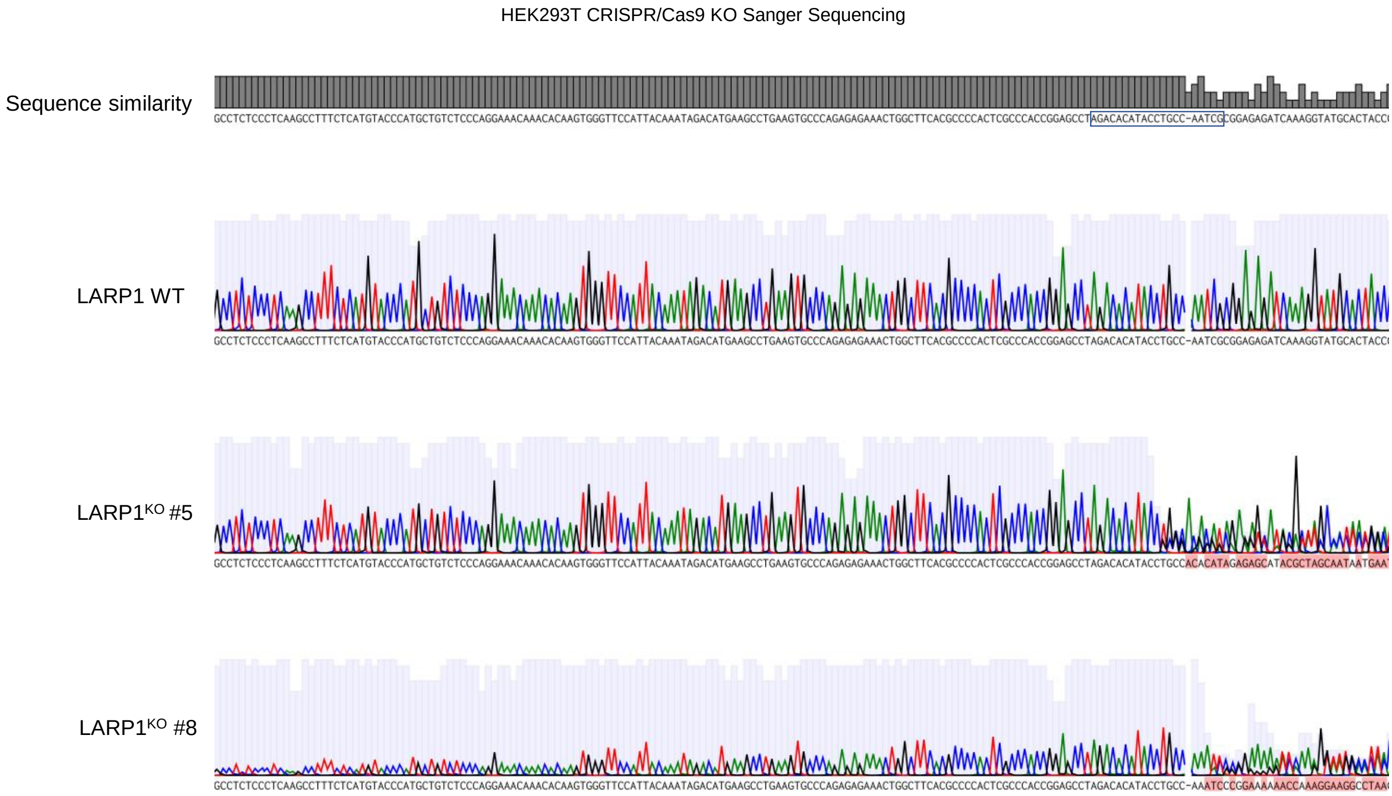


B

HEK293T CRISPR/Cas9 KO Genomic Sequence inferred from RNA-Seq

| Clone | | Genomic Sequence | Mutation |
|---|---|---|---|
| LARP1 Wildtype | | 5'-TAGACACATACCTGCCAATCGCGGAGAGATCAAA-3' | wildtype sequence |
| LARP1 KO #5 | Allele 1 | 5'-TAGACACATACCTGCCA* *CGCGGAGAGATCAAA-3' | * * denotes 2nt (AT) deletion |
| | Allele 2 | 5'-TAGACACATACC* * * * * * * *GCGGAGAGATCAAA-3' | * * * * * * * * denotes 8nt (TGCCAATC) deletion |
| LARP1 KO #8 | Allele 1 | 5'-TAGACACATACCTGCCAA**A**TCGCGGAGAGATCAAA-3' | _ denotes insertion of 1nt (A) |
| | Allele 2 | 5'-TAGACACATACCTGCCAA**AA**TCGCGGAGAGATCAAA-3' | _ denotes insertion of 2nt (AA) |

C

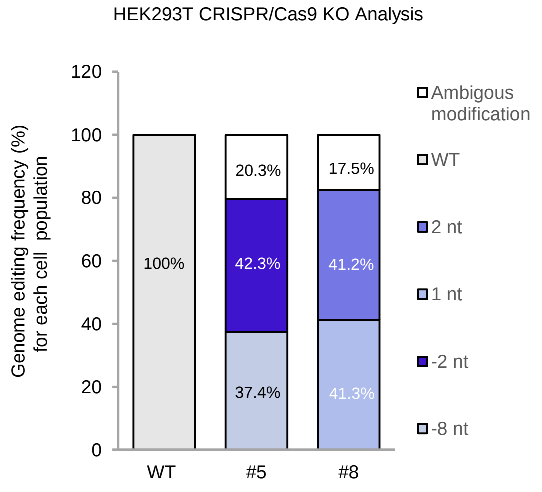


D

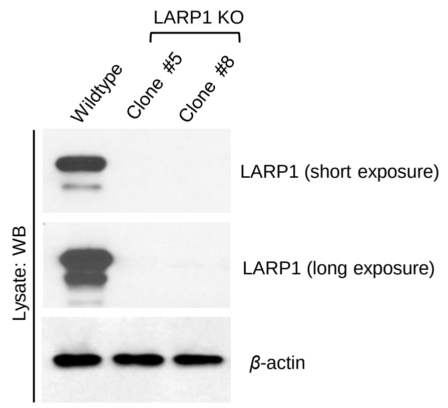


**Supplemental Figure 5.** Genomic editing of human *LARP1* gene locus by CRISPR/Cas9. (A) Sanger sequencing chromatograms of PCR products spanning the sgRNA targeting site (boxed sequence) from unedited (LARP1 WT) and edited clones. (B) RNA sequencing results for LARP1 mRNA isolated from wildtype (parental) or LARP1 CRISPR/Cas9 KO clonal cell lines. (C) Frequency (%) of genome editing events within *LARP1* genomic region for each of the CRISPR/Cas9 cell clones determined by TIDE analysis of Sanger sequencing chromatograms (shown in A). *2nt* denotes 2 nucleotide insertion; *1nt* denotes 1 nucleotide insertion; *-2nt* denotes 2 nucleotide deletion; *-8nt* denotes 8 nucleotide deletion; Sanger sequencing reads that suggested genomic alterations but lack statistical power are shown as *ambiguous modifications*. **(**D) SDS-PAGE/Western blot analysis of HEK293T lysates from wildtype (parental) or LARP1 CRISPR/Cas9 KO clonal cell lines.
